# New proteomic signature in circulating extracellular vesicles from tumor-draining vein of lung adenocarcinomas patients

**DOI:** 10.1101/2024.10.28.620567

**Authors:** Jérémy Tricard, Stéphanie Durand, Amy Gateau, Luc Negroni, Alain Chaunavel, François Bertin, Massimo Conti, Hussein Akil, Fabrice Lalloué

**Author notes:** **Correspondence :** Hussein Akil,; Fabrice Lalloué. These authors contributed equally to this work (Jérémy Tricard, Stéphanie Durand). These authors contributed equally to this work (Hussein Akil, Fabrice Lalloué).

## Abstract

Identification of noninvasive prognostic biomarkers, allowing monitoring of frequently developed relapse in patients with locally advanced non-small cell lung cancer (NSCLC), still of primary importance. Tumor-draining vein (TDV) plasma samples, are known to be enriched in circulating cancer biomarkers compared to samples from peripheral vein (PV). Thus, we thought to investigate the proteomic profile of extracellular vesicles (EVs) from TDV compared to those from PV plasma samples of patients operated for NSCLC.

Purified EVs from TDV and PV plasma samples were characterized for their size distribution and concentration using nanoparticles tracking analysis (NTA). Proteomic profiling of TDV-derived EVs and PV-derived EVs were further done using mass spectrometry (nanoLC-MS/MS) analysis. In parallel, proteomic profile of tumoral and non-tumoral adjacent counterpart tissues from patients with NSCLC were investigated.

Twenty patients with NSCLC, treated by surgery with curative intent, were enrolled in this study. We showed that EVs from TDV plasma samples were significantly smaller than those from PV plasma samples. Interestingly, the concentration of TDV-derived EVs were significantly higher than PV-derived EVs. However, EVs concentration and size were not associated with tumor size or other clinical characteristics. Proteomic profiling showed that 9 of the 10 most overexpressed proteins in EVs from TDV samples compared to those from PV, were associated with lung cancer diagnosis and prognosis. Remarkably, 1 protein (SRPRB) was commonly upregulated in lung tumor tissues (as compared to non-tumoral counterparts) and in TDV-derived EVs (as compared to PV-derived EVs). In contrast, 12 proteins were found to be upregulated in TDV-derived EVs and downregulated in tumor tissues.

In conclusion, all of these identified proteins, carried by EVs from TDV plasma samples, might represent promising novel biomarkers for NSCLC prognosis and predicting recurrences at early stages.

## INTRODUCTION

Non-small cell lung cancer (NSCLC) is the leading cause of cancer deaths worldwide [1], and adenocarcinoma is the most common cancer type (∼80%). Surgery is the corner stone of treatment of early-stage and resectable locally advanced NSCLC [2]. Nevertheless, patients frequently develop local and/or distant relapses despite complete surgical resection. Rate of recurrence is approximately 25% for stage I-II NSCLC [3] and 50-70% for stage III NSCLC, treated with multimodality therapy [3, 4]

Thus, identification of prognostic markers of early relapse after surgery is a major challenge. Indeed, currently used imaging techniques unfortunately are not sensitive enough to detect metastasis at an early stage. Several liquid biopsy biomarkers have been studied to predict resected cancer recurrence, but none are used in current practice for this purpose, including circulating tumor cell (CTC) [5–7], circulating tumor DNA (ctDNA) [8, 9]. Among them, extracellular vesicles (EVs) are nanoscale lipid bilayer vesicles secreted by all cells, acting as key mediators of intercellular communication[10]. The EVs released from tumor cells, including exosomes, are involved in premetastatic niche establishment, tumor progression, angiogenesis, and immune response modulation [10, 11]. Indeed, they are known for their capacity to transport many cellular components, such as mRNA, miRNA, lipids and proteins. Thus, EVs cargo could reflect the characteristics of the cell of origin.

Due to their abundance and stability in biofluids, EVs analysis could be more relevant over conventional liquid biopsy [10–12]. Nevertheless, the use of EVs as biomarkers might overcome some limitations associated with CTCs or ctDNA, especially their very low concentration detected in the blood of patients.

It’s worth mentioning than tumor-derived EVs represent a minor population of circulating EVs released by all cells of the body [13, 14], particularly in non-metastatic stage diseases. To limit this pitfall, many studies indicated that tumor-draining vein (TDV) samples, are enriched for many oncological biomarkers when compared to samples from peripheral blood. This difference could be explained by the dilution effect of tumor-derived EVs in the systemic blood and the immune system degradation [15].

Navarro and colleagues was the only research team which have studied EVs of tumor-draining pulmonary vein in patients operated for NSCLC [16–18]. They showed that patients with smaller pulmonary vein exosome presented high risk of relapse following surgery and worse overall survival [16]. Furthermore, miRNA analysis of EVs isolated from pulmonary vein in the operated patients cohort revealed potential relapse biomarkers [17, 18].

To our knowledge, proteomic analysis of EVs from vascular bed of lung cancer has never been performed and protein cargo of peripheral blood EVs in lung cancer patients revealed some signatures associated with diagnosis [19–22] and prognosis [23, 24].

Considering these research approaches, we hypothesized that blood samples in pulmonary vein are enriched in primary tumor-derived EVs and conducted the first proteomic analysis of EVs from operated lung adenocarcinoma-draining vein, compared with peripheral EVs protein cargo and cancer tissue.

## MATERIALS AND METHODS

### Study design

ExOnSite-Pro (Molecular Profiling of Exosomes in Tumor-draining Vein of Early-staged Lung Cancer, Clinical Trial Register number: NCT04939324, ethical approval the 21^st^ may 2021 by the independent protection committee SUD MEDITERRANEE III – 2021.05.02 bis_21.03.15.45551, FRANCE) is a prospective study performed from June 2021 to June 2024. Written informed consent was obtained for each patient.

We included 20 patients operated on curative-intent lung adenocarcinoma resection at Dupuytren University Hospital of Limoges (Limoges, France). Patients with cancer history, induction therapy, or non-free resection margins were not included.

### Patient samples

After resection, tumoral and non-tumoral tissues were snap frozen in liquid nitrogen and conserved at -80°C. Blood (20 mL) was collected by the nurse anesthesiologist in EDTA tubes, from peripheral vein before the start of surgery. The thoracic surgeon punctured the pulmonary tumor-draining vein (upper pulmonary vein for upper and middle lobe tumors, lower pulmonary vein for lower lobe tumors) before starting resection, to collect 20 mL of blood in EDTA tubes. Plasma obtained by centrifugation was stored at -80 °C.

### Extracellular vesicles (EVs) purification

EVs were purified according to standard protocol described by Théry *et al*., [25]. Briefly, 0.5 mL of plasma was first diluted with an equal volume of PBS (Phosphate-Buffered Saline) and centrifuged at 2000g for 20 minutes (min), the resultant supernatant was then centrifuged at 16 500g for 45 min. Supernatant was further submitted to ultracentrifugation at 120 000g for 2 hours (h) using Optima MAX-XP Beckman coulter ultracentrifuge and MLA-130 rotor. Pellet was then resuspended with PBS and submitted to ultracentrifugation at 120 000g for 2 h. Finally, pellet was resuspended in PBS for NTA (Nanoparticles Tracking Analysis) or flow cytometry analysis.

For proteomic analysis, EVs were purified from plasma using SEC (Size-Exclusion Chromatography) technique. IZON qEV2 Gen2, 35 nm series columns were used following the manufacturer’s instructions. Briefly, 2 mL of plasma were loaded onto the column for each patient and 5 fractions of 2 mL were eluted in PBS. The eluted plasma-EVs fractions were then ultracentrifuged at 120 000g for 2 h. Pellets containing EVs were resuspended with 20 µL cell lysis buffer from Cell Signaling Technology and stored at -80°C.

### Extracellular vesicles (EVs) characterization

#### Nanoparticle Tracking Analysis (NTA)

NTA was performed using NanoSight NS300 (Malvern Panalytical Ltd.) as previously described [26]. Captures and analysis were achieved by using the built-in NanoSight Software NTA3.3.301 (Malvern Panalytical Ltd). The camera level was set at 14, and the detection threshold was fixed at 5 Samples were diluted in PBS to a final volume of 1 mL, and their concentration was adjusted by observing a particles/frame rate of around 50 (30–70 particles/frame). For each measurement, five consecutive 60 seconds (s) videos were recorded under the following conditions: cell temperature 25 °C, syringe speed 22 microL/s (100 a.u.). Particles (EVs) were detected using a 488 nm laser (blue), and a scientific CMOS camera. Among the information given by the software, the following were studied: mean size, mode (i.e., the most represented EVs size population), and particles/mL.

### Flow cytometry analysis

EVs resuspended in PBS were first immunocaptured using magnetic beads coated with anti-CD81 antibody (Thermofisher, Dynabeads Cat N° 10616D) according to the manufacturer’s instructions. The complexes formed by the beads-bound EVs were then stained with anti-CD81 APC antibody (Biolegend, clone 5A6) and signal was detected using Cytoflex flow cytometry (Beckman Coulter). Data were analyzed using Kaluza software.

### Proteomic analysis Sample preparation

Total protein lysates were extracted from tissue and purified EVs samples using cell lysis buffer (20 mM Tris-HCl pH 7.5, 150 mM NaCl, 1 mM Na2EDTA, 1 mM EGTA, 1% Triton, 2.5 mM sodium pyrophosphate, 1 mM beta-glycerophosphate, 1 mM Na3VO4, 1 µg/ml leupeptin, 1 mM PMSF. After 30 min of incubation on ice, samples were briefly sonicated and then centrifuged at 16 000g for 10 min. Total protein concentrations were determined using Pierce BCA protein assay kit (Thermo Fisher).

### Mass spectrometry analysis

#### Liquid digestion

After a protein precipitation with TCA overnight at 4°C, pellets were washed twice with 1 mL cold acetone, dried then dissolved in 1 M urea in 0.1 mM Tris-HCl pH 8.5 for reduction (10 mM DTT, 30 min, 56°C), and alkylation (20 mM iodoacetamide, 30 min, 25°C). Digestion was performed by two addition of trypsin (T_O_ and T_2h_) for overnight digestion at 37°C. After addition of TFA (0.2% final), the sample were ready for MS analysis.

#### Mass spectrometry

nanoLC-MS/MS was performed with the Ultimate 3000 nano-RSLC coupled in-line with a Exploris 480 quadrupole-orbitrap via a nano-electrospray ionization source and the FAIMS pro interface (Thermo Scientific, San Jose California). Tryptic peptides (1 µl) were loaded on the preconcentration cartridge (C18 PepMap100 trap-column 300µm x 1 mm) for 1 min. at 15 µL/min with 2% ACN, 0.1% FA in H_2_O and separated on analytical column (C18 PepMap 75 µm ID x 15 cm, Thermo Fisher Scientific) with a 40 min. gradient from 8% to 25% buffer B (A: 0.1% FA in H_2_O; B: 0.1% FA in 80% ACN, 450 nl/min, 45°C) followed by a regeneration step at 90% B and a equilibration to 8% B. Total chromatography time was 60 min. The mass spectrometer was operated in positive ionization mode and Data-Dependent Acquisition (DDA) with 2 cycles of FAIMS compensation voltages (-45V and -55 V for 1.2 and 0.8 sec, respectively). The two FAIMS-DDA cycle consisted of one survey scans (350-1200 m/z, 60,000 FWHM) followed by MS² spectra (HCD; 30% normalized energy; 2 m/z window; 22,500 FWMH). The Normalized AGC were 300% and 100% for MS1 and MS², respectively, with a maximum injection time set to 50 ms for both scan modes. Unassigned and single charged states were rejected. Exclusion duration was set for 40 s with mass width was ± 10 ppm.

#### MS data processing

Proteins were identified with Proteome Discoverer 2.5 software (Thermo Fisher Scientific) and *Homo Sapiens* proteome database (Swissprot, reviewed, release 2024-03-08, 20597 sequences). Precursor and fragment mass tolerances were set at 7 ppm and 0.05 Da, respectively, and up to 2 missed cleavages were allowed. Oxidation (M, +15.9949) was set as variable modification, and Carbamidomethylation (C, +57.021) as fixed modification. Peptides were filtered with a false discovery rate (FDR) at 1%. Proteins were quantified with a minimum of 1 unique peptide based on the XIC (sum of the Extracted Ion Chromatogram). All raw LC-MS/MS data have been deposited to the ProteomeXchange via the PRIDE database.

### Bioinformatic treatment of proteomic data

The quantification values were exported in R environment for statistical analysis, using DEP package [27] involving data filtering, variance normalization and imputation of missing values before differential analysis using linear model to identify differentially enriched / expressed proteins [27]. We first filter out proteins associated to a Peptide Spectrum Match (PSM) inferior or equal to 3, and second, we filter proteins that contain too many missing values. Only proteins detected in at least 70% of samples will be retained for subsequent analysis. Then, the data is background corrected and normalized by variance stabilizing transformation and the remaining missing values in the dataset were imputed using k-nearest neighbor method. Protein-wise linear models combined with empirical Bayes statistics are used for the differential expression analysis with comparison of tumor compared to normal from proteome data of tissue samples (19 T *versus* 20 NT) and comparison of tumor-draining vein compared to peripheral vein from proteome data of EVs samples (19 TDV *versus* 14 PV). Thresholds for selection of differentially detected protein are: absolute log_2_ fold change superior or equal to 1 and adjusted p value (corrected by Benjamini-Hochberg multiple testing method) inferior or equal to 0.05. The results of differential analysis were visualized using volcano plot (EnhancedVolcano package) [28] and hierarchical clustering analysis (ComplexHeatmap package) [29]. Principal component analysis was performed using FactoMineR and factoextra packages.

### Data integration into biological networks

To obtain biological information about this set of deregulated genes in EVs from TDV as compared to EVs from PV, we searched for direct (physical) and indirect (functionally associated with no direct interaction) associations between 369 proteins identified by proteome analysis in the STRING (Search Tool for the Retrieval of Interacting Genes/Proteins) database (version 12.0) [30]. Briefly, interactions in STRING are derived from multiple sources (experimental/biochemical experiments, curated databases, genomic context prediction, co-expression and automated text mining). Confidence level of edge, i.e. interaction between 2 nodes or protein, is computed in a combined score (ranging from 0 to 1). Interactions and proteome data were imported and used for network generation in the Cytoscape environment [31] (version 3.10). Topological analysis was performed with Cytohubba [32] to identify nodes parameters, including degree and bottleneck.

### Statistical analysis

The Mann-Whitney U test was used de determine statistical significances of NTA analyses.

## RESULTS

### Characterization of EVs purified from tumor-draining vein and peripheral vein of lung cancer patients

The schematic workflow illustrating the various stages of our study from collection of blood samples at tumor resection to *in silico* analysis of EV-derived proteome derived from EVs are shown in figure 1. Patients’ characteristics and operative data are detailed in table 1. All margins resections were free. Height patients (40%) experienced adjuvant therapy (6 treated by cisplatin-based chemotherapy, 1 by chemotherapy and osimertinib and 1 by osimertinib). After purification, EVs derived from tumor-draining vein (TDV) or peripheral vein (PV) of lung cancer patients were characterized for their size distribution by nanoparticle tracking analysis (NTA). As shown in figure 2A, EVs from both TDV samples and PV samples are between 50 and 150 nm in size, consisting with the characteristic size of small EVs. In parallel, flow cytometry analysis (Figure 2B) confirmed the presence of CD81 tetraspanin protein, usually used as EVs marker. These results confirmed the successful purification of EVs from TDV and PV plasma samples.

**Figure 1:**
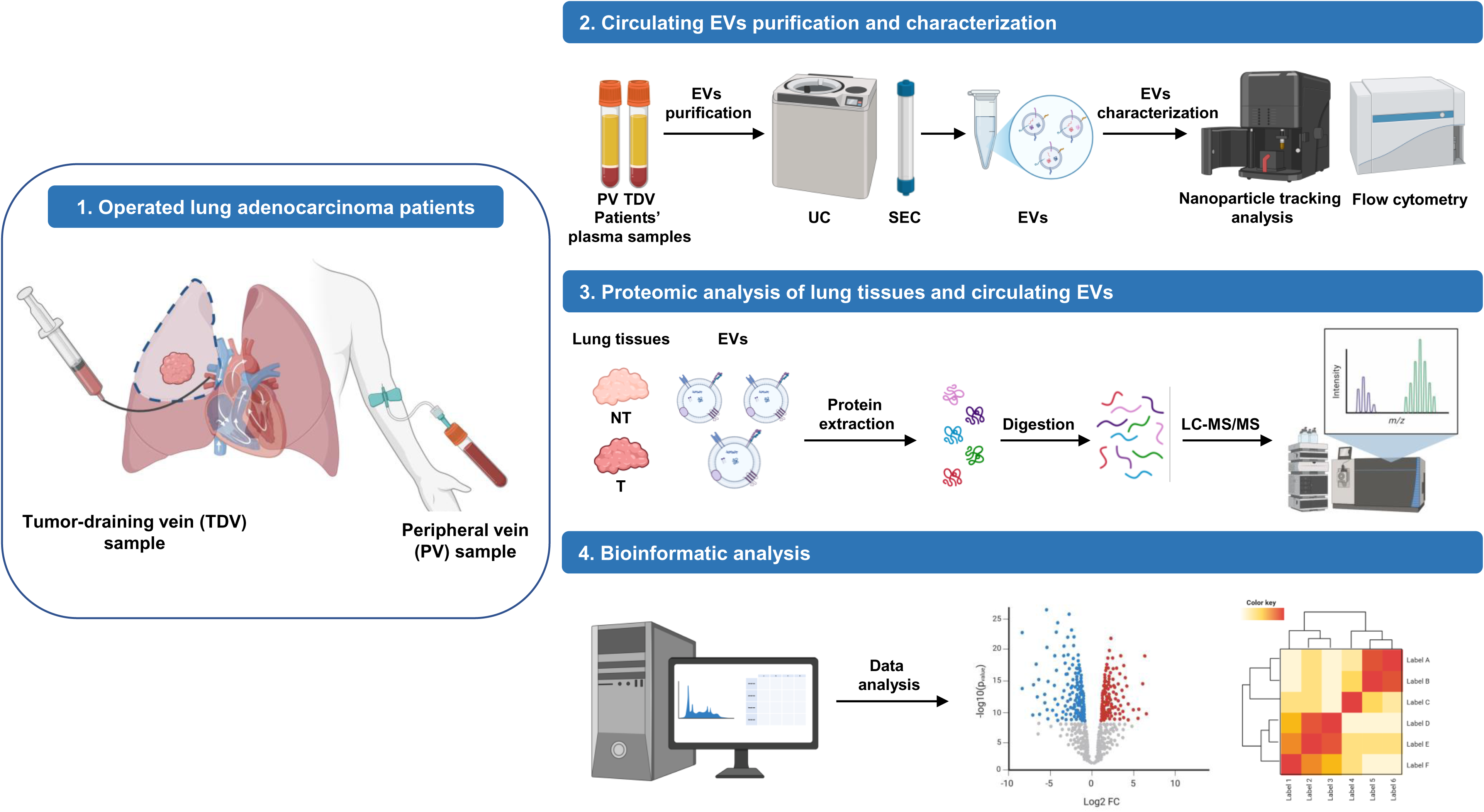
schematic representation of the study workflow. EVs: extracellular vesicles; UC: ultracentrifugation; SEC: size exclusion chromatography; NT: non-tumoral tissues; T: tumoral tissues; LC-MS/MS: liquid chromatography-tandem mass spectrometry. Figure created with BioRender.com

**Figure 2:**
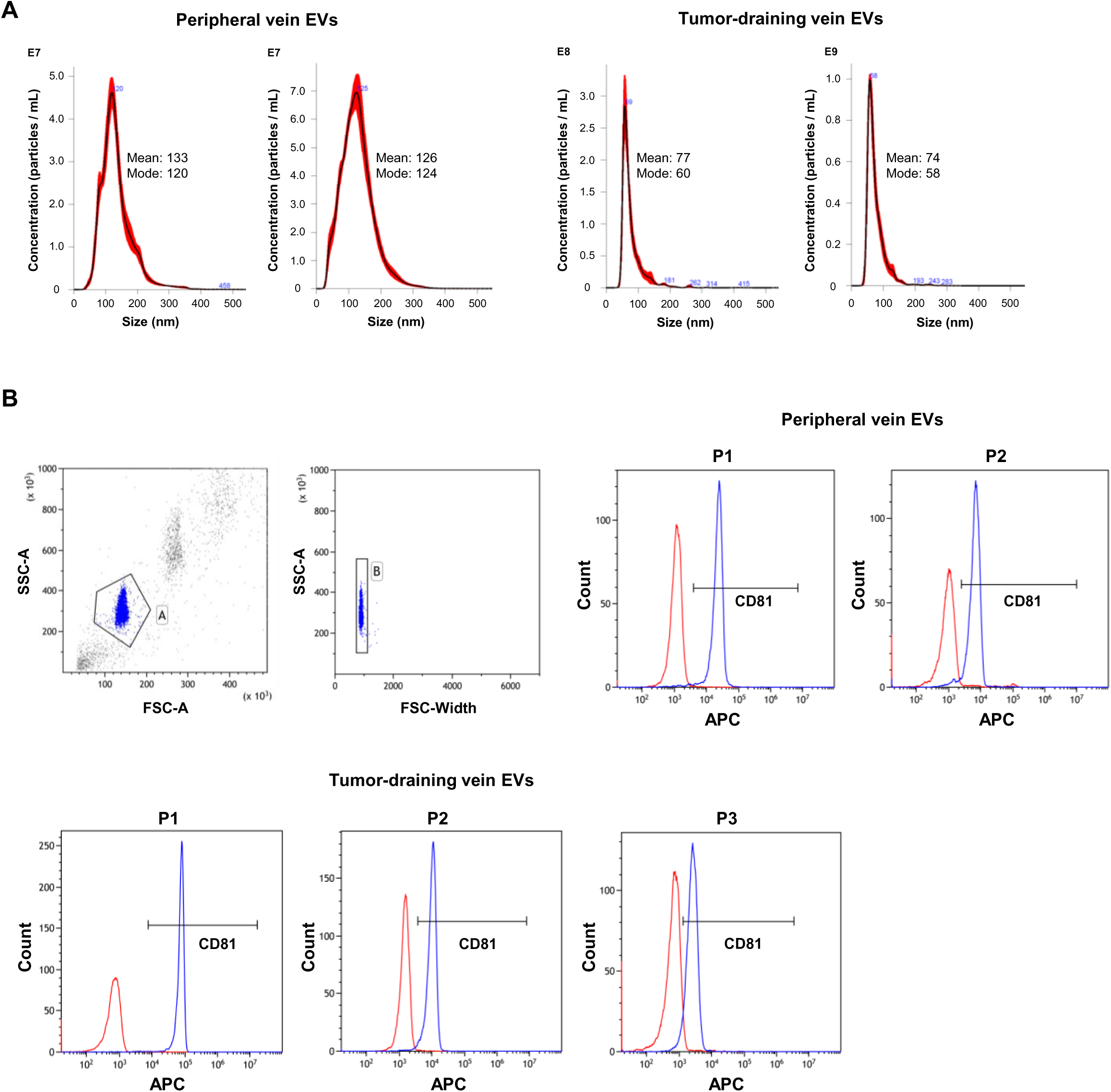
Characterization of circulating extracellular vesicles (EVs) purified from peripheral vein (PV) or tumor-draining vein (TDV) plasma samples. (A) Representative nanoparticle tracking analysis (NTA) of circulating EVs purified from peripheral blood (left) or tumor-draining blood (right) of 2 lung cancer patients. Black line represents the mean value and the red shaded area represents the standard error of 5 recordings. (B) Flow cytometry analysis of CD81 on circulating EVs purified from peripheral vein (N=2) or tumor-draining blood (N=3) of lung cancer patients. Circulating EVs were incubated with anti-CD81 magnetic beads and counter stained with anti-CD81 APC or isotype control conjugated to APC. The gating strategy (left top) is presented in the upper left part. Red histograms correspond to isotype control and blue histograms correspond to CD81 staining.

**Table 1.**
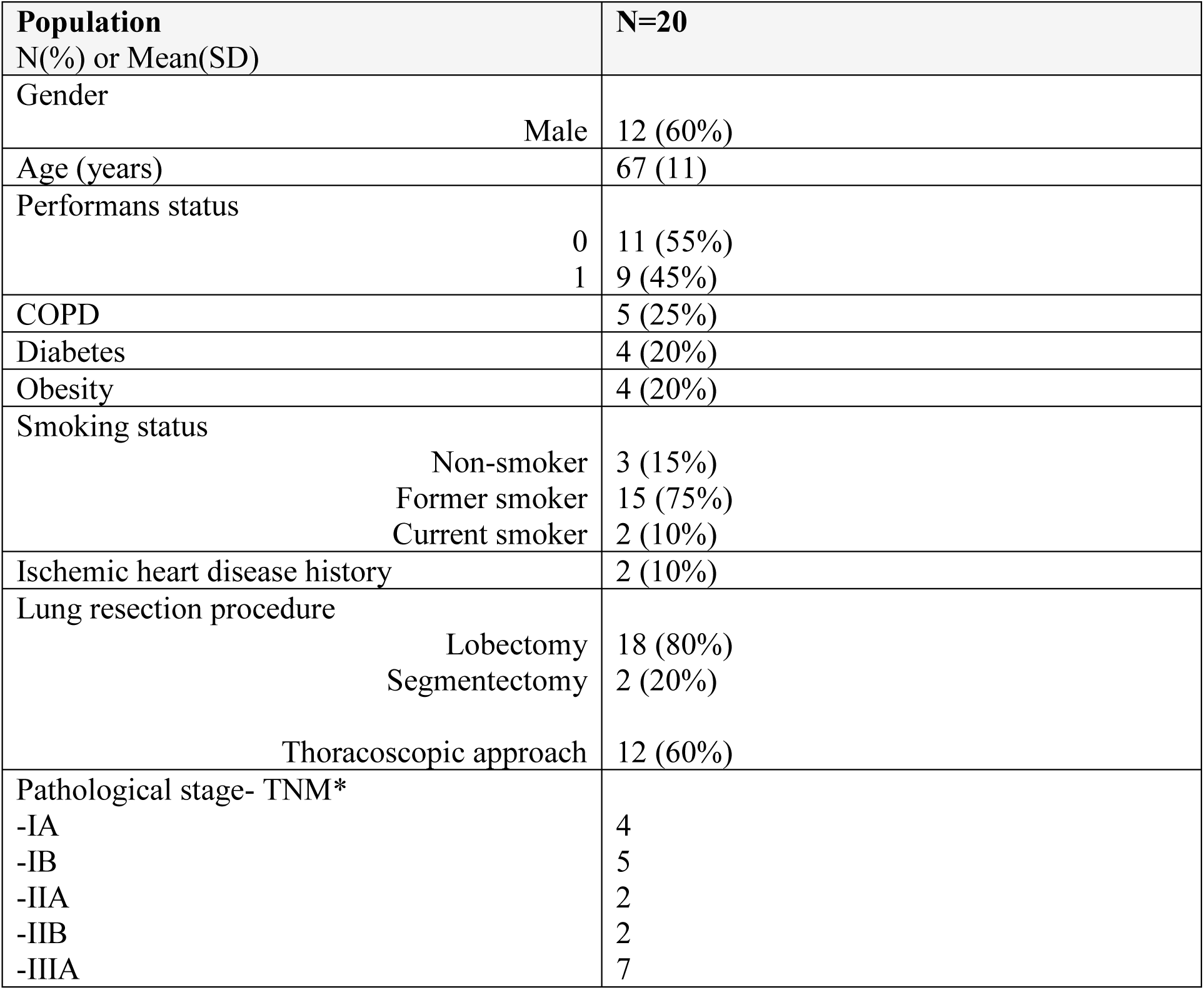
Study population. N(%): effective (percentage) ; SD: standard deviation. *according to the 8th edition of the classification.

### EVs are smaller and more abundant in TDV plasma samples compared to those in PV plasma samples

Based on NTA analysis (Figure 3A), EVs concentration was significantly higher in TDV plasma samples compared to PV plasma samples (17.0 10^9^ vs 4.4 10^9^ particles/mL respectively). In addition, EVs in TDV plasma samples were smaller than EVs in PV plasma samples (respectively, mean size: 98.7+/-22.9 nm vs 120.6+/-23.4 nm; mode size: 74.7+/-17.5 vs 97.2+/-25.2 nm) (Figure 3A). Interestingly, analysis according to three size groups (< 100 nm, 100-150 nm and > 150 nm) showed that TDV plasma samples were significantly enriched in three groups of EVs compared to those in PV plasma samples. Furthermore, EVs with a size < 100 nm were significantly enriched in TDV plasma samples when compared to TDV EVs group with a size ranging between 100 and 150 nm (Figure 3B). Although EVs in TDV plasma samples, with a size < 100 nm, tend to be higher than those with a size > 150 nm, the observed difference is not significant.

**Figure 3:**
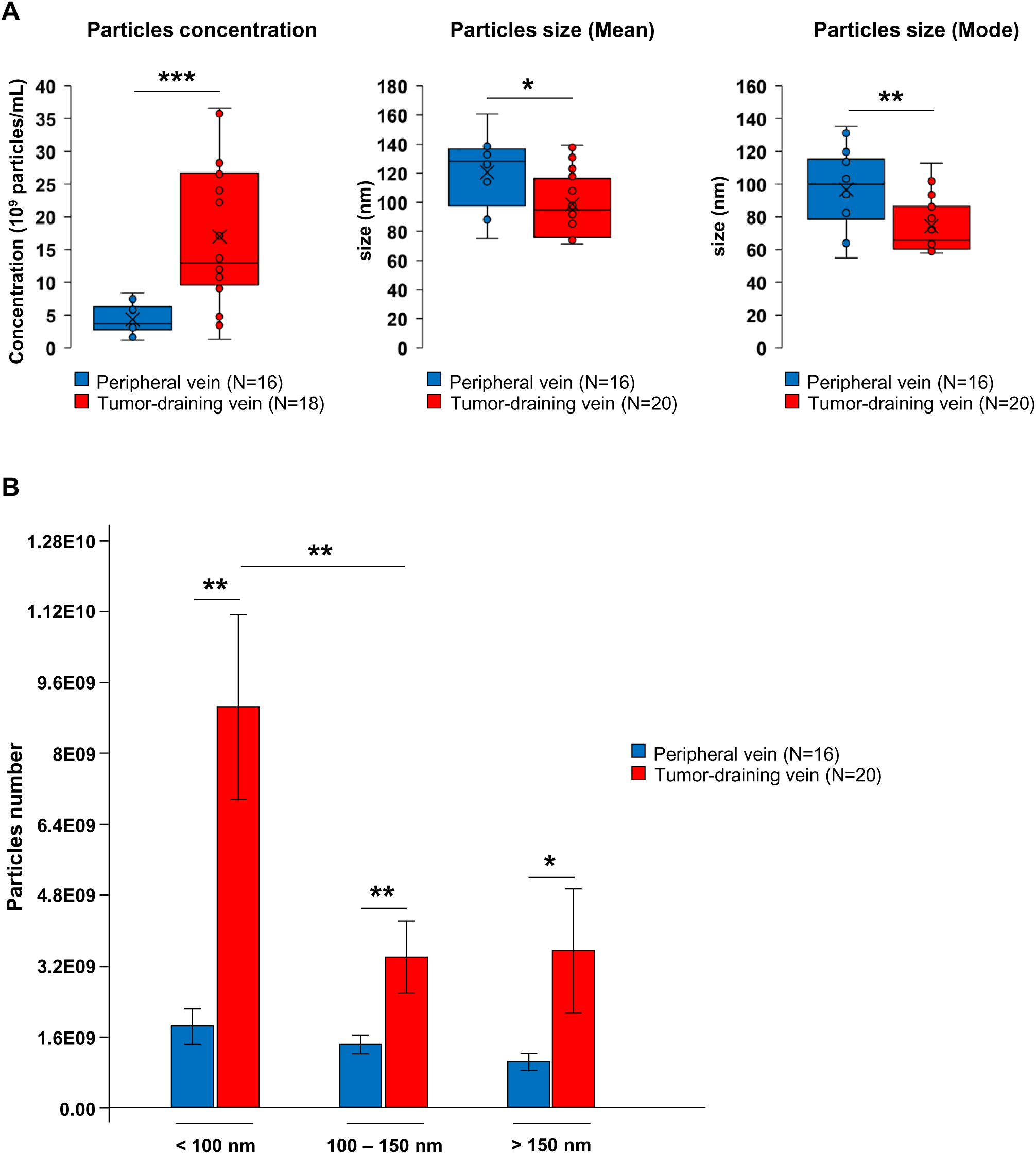
EVs are smaller and more abundant in TDV compared to PV plasma samples. (A) NTA data are presented as box plots for particles concentration (left), mean size (middle) and mode size (right) obtained from peripheral vein or tumor-draining vein. (B) Histograms representing EVs size levels purified from peripheral vein or tumor-draining vein. EVs were divided into three groups: below 100 nm (< 100 nm), between 100 and 150 nm (100 – 150 nm) and higher than 150 nm size (> 150 nm). (* *p* < 0.05, ** *p* < 0.01, *** *p* < 0.001).

EVs concentration and size were not associated with tumor size or other clinical characteristics. In our cohort, four patients (20%) relapsed after surgery, 3 with distant metastases. All of them presented pIIIA stage-cancer. No patient died during follow-up. Risk factors for cancer recurrence was operative blood loss (median 105 ml vs 40 ml, p=0.047), tumor size (72 +/-21 mm vs 31+/-14 mm, p=0.024) and pN+ status (p=0.013).

EVs concentration, median size and median mode in PV plasma samples were similar when comparing patients who experienced relapse and those who did not (respectively, 3.23 vs 3.77, p=0.7; 137 vs 126, p=0.19; 107 vs 94.3, p= 0.19). EVs concentration, mean size and mean mode in TDV samples were also similar between patients who experienced relapse and those who did not (respectively, 18.0 vs 16.7, p=0.78; 96.8 vs 99.2, p=0.84; 72.0 vs 75.4, p=0.89).

### Mass spectrometry analysis revealed poor prognosis-associated proteins in lung adenocarcinoma tissues

Among the 3878 proteins characterized by liquid Chromatography tandem MS (LC-MS/MS) with a Peptid Spectrum Match (PSM) strictly superior to 3, 2758 proteins were quantified in all patients (19 tumor and 20 paired non-tumoral tissues) and 3553 proteins were expressed in more than 70% of patients. Principal component analysis (PCA) analysis demonstrated a marked segregation between tumor (T) and healthy tissue (NT) proteomes suggesting that tumor progression and growth altered proteins expression profile (Figure 4A). Among 3553 proteins, 148 proteins (31,75 %) were significantly increased and 318 proteins (68,25 %) were decreased compared non-tumoral tissues (Supplementary table 1). 148 and 318 proteins had a more than 2-fold increase or decrease, respectively. Some of these proteins (72) are overexpressed up to 4-fold increase or decrease more than others (Figure 4B). To investigate what protein expression are influenced by tumor development, we also conducted clustering analyses on the tumor and normal tissues. We showed three distinct clusters. The first cluster (1) mainly includes non-tumor tissues (except for 1 tumor tissue) whereas the second (2) only gathered tumor. The third can be considered as an intermediate subgroup mixing non-tumor and tumor samples (Figure 4C). This distinction in 3 subgroups highlighted three distinct proteomes. Indeed cluster 1 and 2 are complete opposites since proteins overexpressed in cluster 1 are downregulated in cluster 2 and reciprocally confirming the clear changes of proteomes between tumor and non-tumor tissues previously described in PCA. This result also enables to identify elevated fold-change overexpressed proteins in tumor that are involved in tumor progression and development in different cancer subtype such as PES1, TMEM165. Among them, several proteins (GOLM1, TUBB3, S100β, CYP1B1, UGT1A6) are already known to be involved in lung cancer progression, metastasis or resistance to treatment. The different clusters were then compared with proteins derived from EVs in order to identify proteins carried by tumor cells-derived EVs and their significance in lung cancer progression.

**Figure 4.**
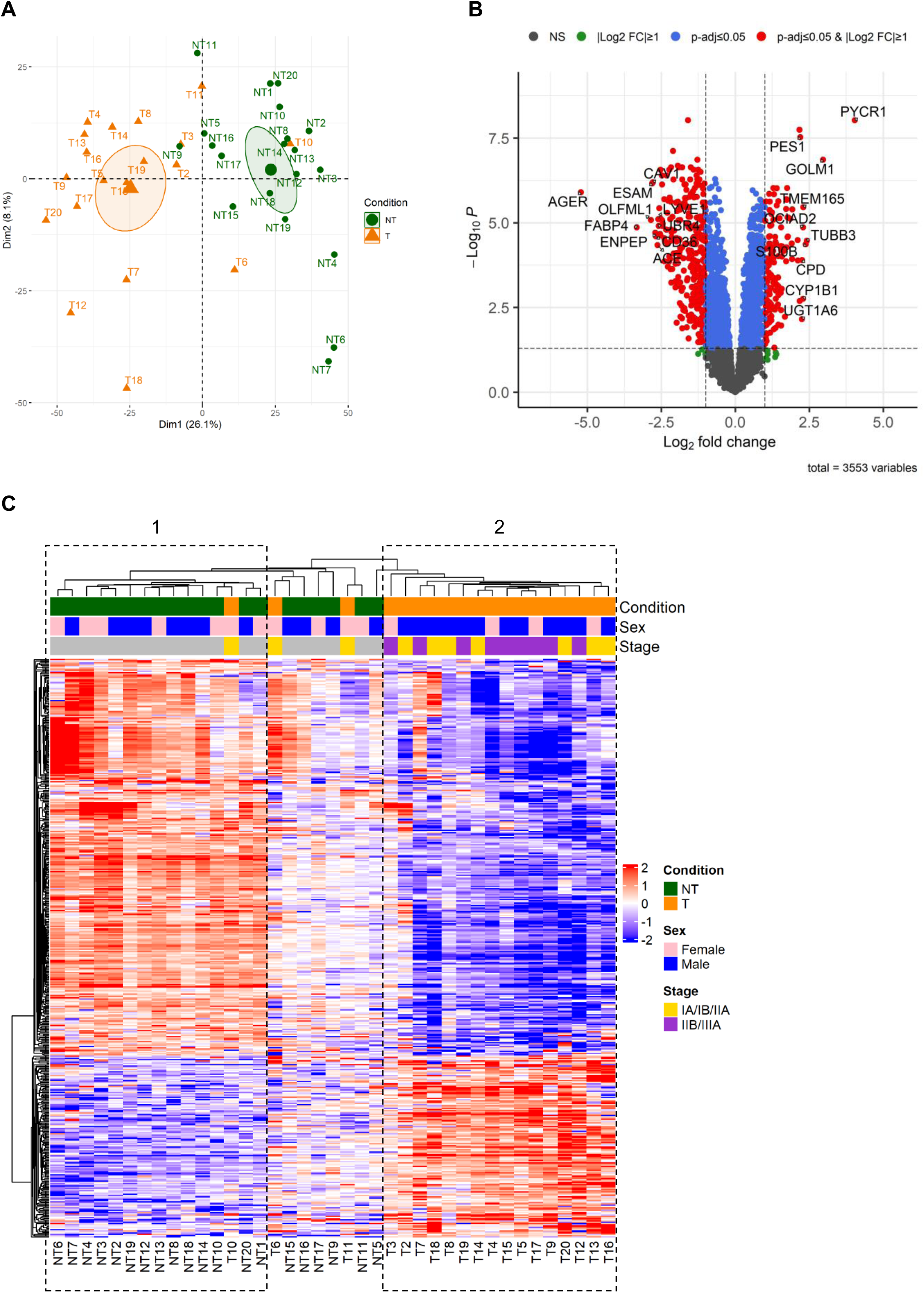
Proteomic profiling of lung adenocarcinoma compared to paired normal tissues. (A) Principal component analysis of proteomic (3553 proteins) data in 39 lung tissue samples (19 tumor and 20 paired normal tissues) reveals a clear distinction between tumor (T, orange triangles) and normal (NT, green dots) lung tissues. The absence of overlap between the confidence ellipses of the barycenter of each group of individuals (*i.e* tumor and normal tissues) confirms their distinct proteomic profiles. (B)Volcano plot resulting of differential analysis between tumor (T) and paired normal tissues (NT). Highlighted points (red) represent 466 proteins showing a significant level change between T and NT with an absolute fold change superior to 2 and an adjusted p value inferior to 0.05 (commonly used threshold in *omic* analysis to identify differentially expressed genes). Dotted lines represents applied thresholds: |(log_2_ FC| ≥ 1 and –log_10_ p.adjusted ≥ 1.3. Among 466 proteins, 148 were up-regulated and 318 down-regulated in tumor as compared to NT. Gene Symbol corresponding to 10 most up- and down-deregulated proteins was indicated. (C) Heatmap of 466 differentially detected proteins in lung tumors (T) as compared paired normal tissue (NT). Protein intensities were log2 transformed and median-centered and are displayed as color gradient from blue to red, reflecting low to high protein level in tissue, respectively. Proteins (rows) and tissue samples (columns) are hierarchically clustered using Pearson correlation distance and average linkage method. Samples characteristics were indicated at the top, with condition (normal tissue: n= 20, green ; tumor: n=19, orange), sex distribution (male: n=12, blue ; female: n=8, pink) and stage (early stage composed IA, IB and IIA pTNM staging: n=10, yellow ; intermediate/late stage represented by IIB and IIIA pTNM staging: n=9, violet).

### Differential protein expression in EVs from tumor draining-vein (TDV) *versus* EVs from peripheral vein (PV)

Among the 1890 proteins characterized by liquid Chromatography tandem MS (LC-MS/MS) in EVs cargo with a Peptid Spectrum Match (PSM) strictly superior to 3, 761 proteins were quantified in all patients (19 patients for EVs from TDV and 14 patients for EVs from PV) and 1410 proteins were expressed in more than 70% of patients. In addition, 87.3% of proteins detected in EVs in our study have already been associated with vesicles in subsequent studies: 78% were retrieved in VesiclePedia and Exocarta databases and 9.3% of proteins were retrieved in at least one of the 2. Interestingly, 12.7% of proteins (179) are new proteins associated to extracellular vesicles (Supplementary figure 1). EVs from TDV seemed to present a protein expression profile different from that of EVs from PV and, PCA analysis showed that the individuals are fairly well separated on PCA (Figure 5A). Among 1410 proteins, 176 proteins (12.5%) were significantly increased and 193 proteins (13.7%) were decreased compared to EVs purified from PV (supplementary table 2). 176 and 193 proteins had a more than 2-fold increase or decrease, respectively. Some of these proteins (94) are overexpressed up to 4-fold increase or decrease more than others (Figure 5B). We conducted clustering analyses of protein expression on EVs from TDV and EVs from PV and showed 2 clusters. The first cluster (1) included a majority of PV-derived EVs (13 *versus* 4 EVs from TDV) whereas the second (2) is composed of only TDV EVs excepting one sample (Figure 5C). Indeed, we identified a specific proteome profile present in EVs from TDV compared to those from PV. These results also allowed to identify specific clusters of overexpressed proteins in EVs from TDV as compared to those from PV. Among these identified proteins, MUC5B and ATP1A are associated with aggressiveness and prognosis of lung cancer and CCT6A, IGVHV5-51, UNC45A which are associated with cancer subtypes.

**Figure 5.**
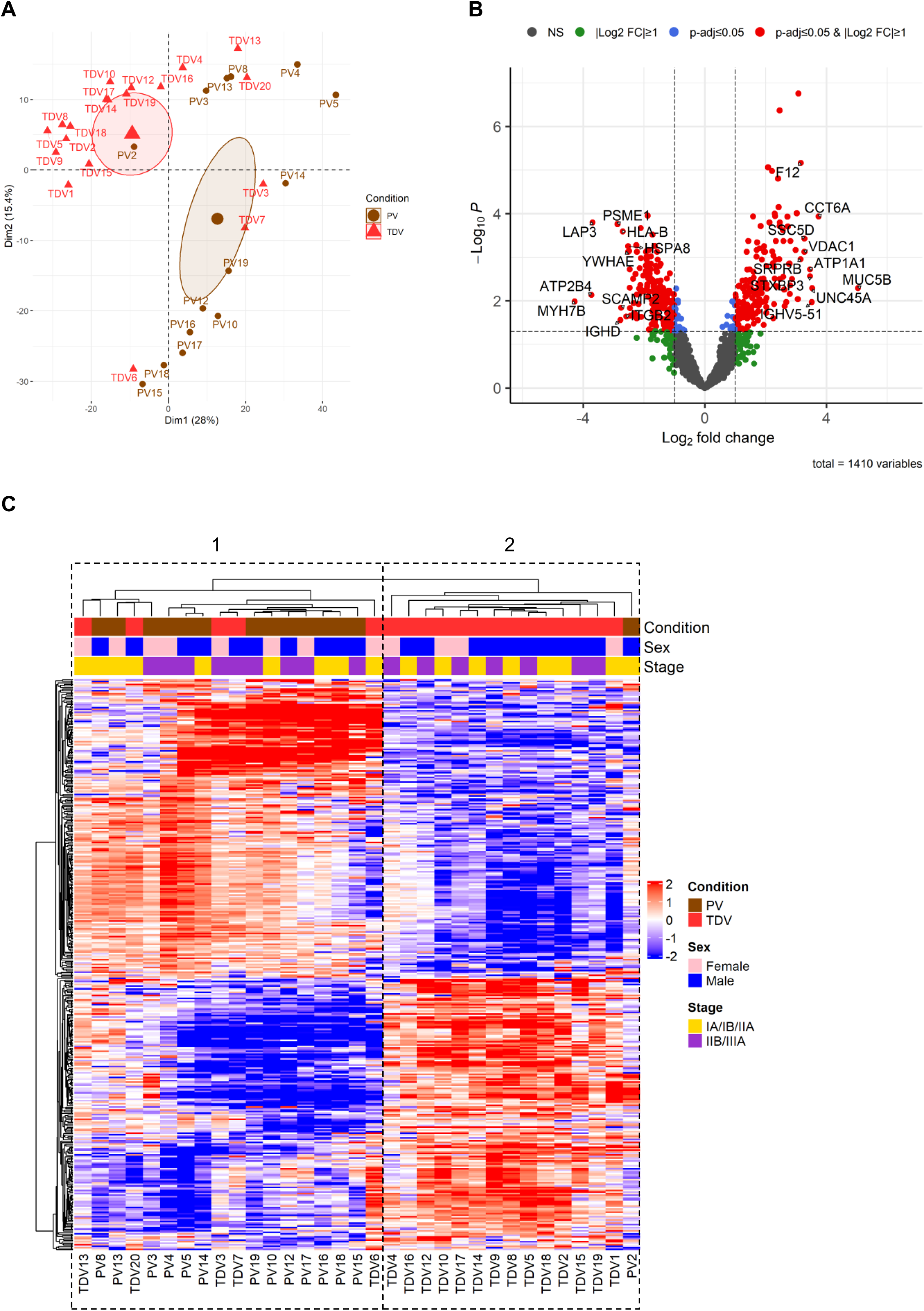
Proteomic profiling of EVs from TDV compared to EVs PV plasma samples. (A) Principal component analysis of proteomic (1410 proteins) data in 33 extracellular vesicles samples purified from Tumor-draining vein (n=19) and peripheral vein (n=14) of lung adenocarcinoma patients reveals a good distinction between EVs from TDV (red triangles) and EVs from PV (brown dots). The absence of overlap between the confidence ellipses of the barycenter of each group of individuals (*i.e* EVs from TDV and PV) confirms their distinct proteomic profiles. (B)Volcano plot resulting of differential analysis between EVs from Tumor-draining vein (TDV) and EVs from Peripheral vein (PV) of patients with lung adenocarcinoma. Highlighted points (red) represent 369 proteins showing a significant level change between EVs from TDV and PV with an absolute fold change superior to 2 and an adjusted p value inferior to 0.05 (commonly used threshold in *omic* analysis to identify differentially expressed genes). Dotted lines represents applied thresholds: |(log_2_ FC| ≥ 1 and –log_10_ p.adjusted ≥ 1.3. Among 369 proteins, 176 were up-regulated and 193 down-regulated in EVs from TDV as compared to EVs from PV. Gene Symbol corresponding to 10 most up- and down-deregulated proteins was indicated. (C) Heatmap of 369 differentially detected proteins in EVs from Tumor-draining vein (TDV) as compared to EVs from Peripheral vein (PV). Protein intensities were log2 transformed and median-centered and are displayed as color gradient from blue to red, reflecting low to high protein level in EVs, respectively. Proteins (rows) and tissue samples (columns) are hierarchically clustered using Pearson correlation distance and average linkage method. Samples characteristics were indicated at the top, with condition (EVs from TDV: n= 19, red ; EVs from PV: n=14, brown) and the gender distribution (male: blue, female: pink) or stage (early stage composed IA, IB and IIA pTNM staging: yellow, intermediate/late stage represented by IIB and IIIA pTNM staging: violet) among these two groups. For TDV group, sex ratio is 12 males/7 females and stage repartition is 10 early stages/9 intermediate-late stages. For PV group, sex ratio is 9 males/5 females and stage repartition is 7 early stages/7 intermediate-late stages.

In order to identify recurrence-associated proteins, we conduct a differential analysis of EVs derived from TDV and those from PV, according to recurrence (supplementary figure 2). For that, hierarchical clustering analysis was conducted from 80 differentially detected protein in EVs from TDV of recurrent patients as compared to no recurrent patients (supplementary figure 2A) and from 92 differentially detected protein in EVs from PV of recurrent patients as compared to no recurrent patients (supplementary figure 2B). Interestingly, despite the low rate of recurrence in our cohort (4/20 patients), recurrent patients were clustered together highlighting a distinct pattern of protein expression in EVs from TDV and PV plasma samples compared to those from non-recurrent patients.

### Comparison of proteome profiling of TDV-derived EVs and tumoral lung tissues

After intersecting the differentially expressed proteins (DEP) of the four conditions [upregulated (148) and downregulated (318) proteins in tumors *versus* non-tumoral tissues with upregulated ((176) and downregulated (193) proteins in TDV-derived EVs *versus* PV-derived EVs], a total of 26 proteins were deregulated and common to EVs from TDV and tumors. One protein (SRPRB) was commonly upregulated in TDV-derived EVs and in lung tumor tissues (figure 6A). Incontrast,12 proteins (AHNAK, APOA2, APOB, APOC1, COL6A1, CSRP1, HLA-E, ITIH2, MCEMP1, PCYOX1, PGLYRP2, SERPING1) were found at the intersection between upregulated proteins from TDV-derived EVs and downregulated proteins in lung tumor tissues (Figure 6A). Furthermore, these data showed that the majority of proteins (46% of deregulated proteins) common to EVs and tumors are downregulated in tumors and instead upregulated in EVs. The overexpression of these proteins in TDV-derived EVs suggests that tumor cells might load these proteins in EVs to prevent their inhibiting function on tumor growth. To further investigate the abundance of tumor-upregulated proteins in EVs, we performed a scatterplot of proteins abundance among 1410 proteins identified in both type of EVs. We determine upregulated proteins in tumor vs normal tissues in order to highlight potential circulating biomarkers of lung adenocarcinoma. Indeed, 128 proteins showed a significant deregulation in lung tumoral tissue (T) when compared to non-tumoral tissues (NT). Eleven proteins were upregulated and 117 downregulated in tumor *versus* non-tumoral tissues (Figure 6B).

**Figure 6.**
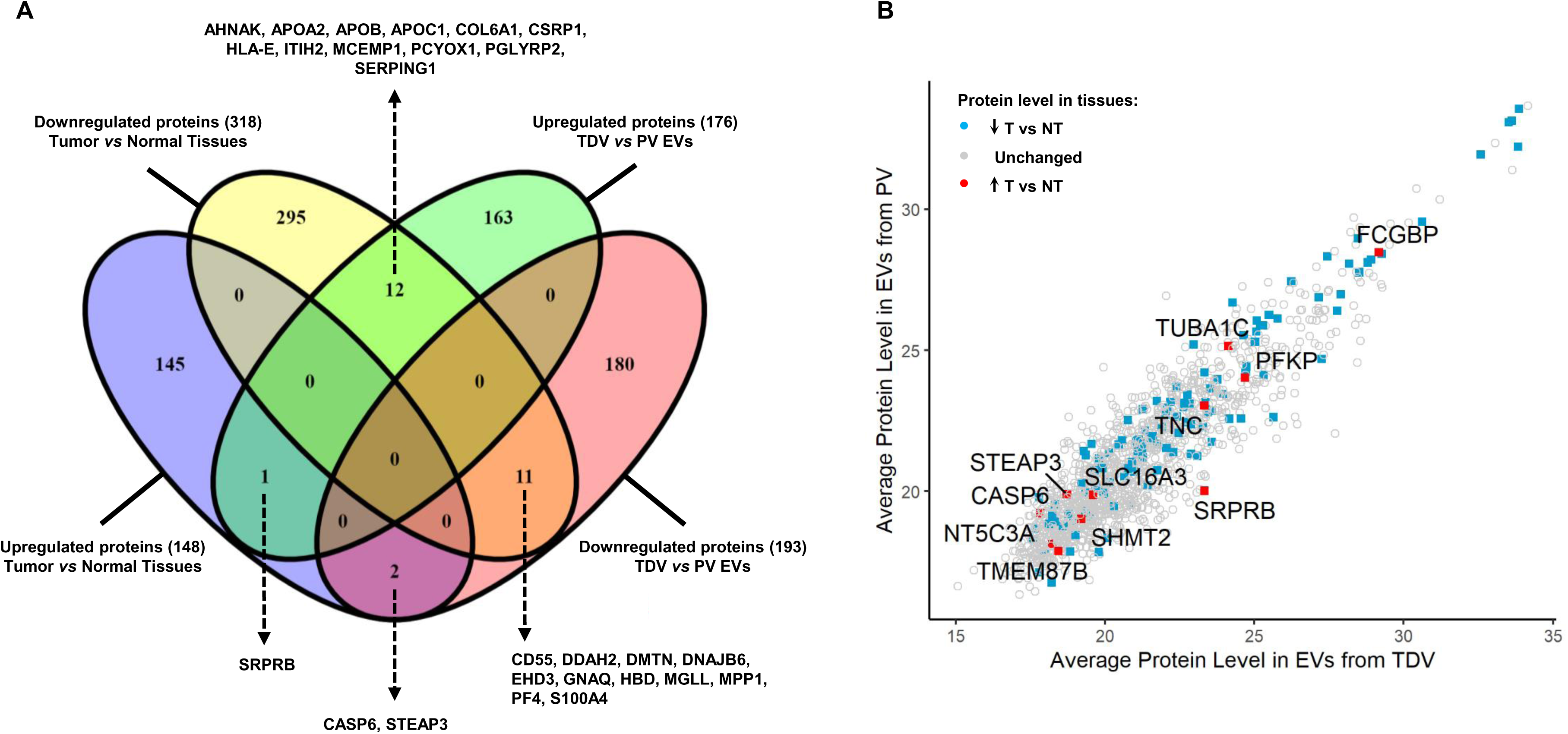
Comparison of proteome profiling of lung tumoral tissues and EVs from TDV plasma samples. (A) Venn diagram between lists of deregulated proteins in lung tumor compared to normal tissue (148 upregulated and 318 downregulated) and lists of deregulated proteins in EVs from Tumor-draining vein (TDV) compared to EVs from peripheral vein (PV) (176 upregulated and 193 downregulated). Gene symbol of commonly deregulated proteins was indicated. (B) Abundance estimation of 1410 proteins-detected in EVs (from TDV and PV) by plotting average protein level in EVs from TDV (n=19) and average protein level in EVs from PV (n=14). Quantification value of protein identified by LC-MS/MS was preliminary normalized, imputed for missing values and log-2 transformed. Among 1410 proteins detected in EVs, 128 show a significant deregulation in lung tumoral tissue (T) as compared to normal tissue (NT) (absolute fold change superior to 2 and adjusted p-value inferior to 0.05): 11 upregulated (indicated by red square) and 117 downregulated (indicated by blue square) in T *vs* NT. Only Gene symbols of upregulated proteins in tumor vs normal tissue were indicated, to highlight these potential circulating biomarkers of lung adenocarcinoma.

### Functional analysis and network building from differentially detected proteins in TDV-derived EVs compared to those from PV samples

The analyses of GO function, KEGG pathways and Reactome Gene Sets indicated that the majority of proteins enriched from GO bound to cellular components correspond to extracellular exosome suggesting that all characterized proteins belong to the exosome cargo (Figure 7A). The top 5 proteins enriched in biological process (BP) are involved in protein transport, regulation of immune system and remodeling of actin cytoskeleton whereas its molecular function are closely linked to enzyme regulation and binding or binding to carbohydrate derivative. Regarding Kegg pathway, the enrichment of at least 50 proteins are observed in signaling by Rho GTPases which are already implicated in mechano-signaling and might regulate osimertinib resistance and brain metastasis in lung cancer [33].

**Figure 7.**
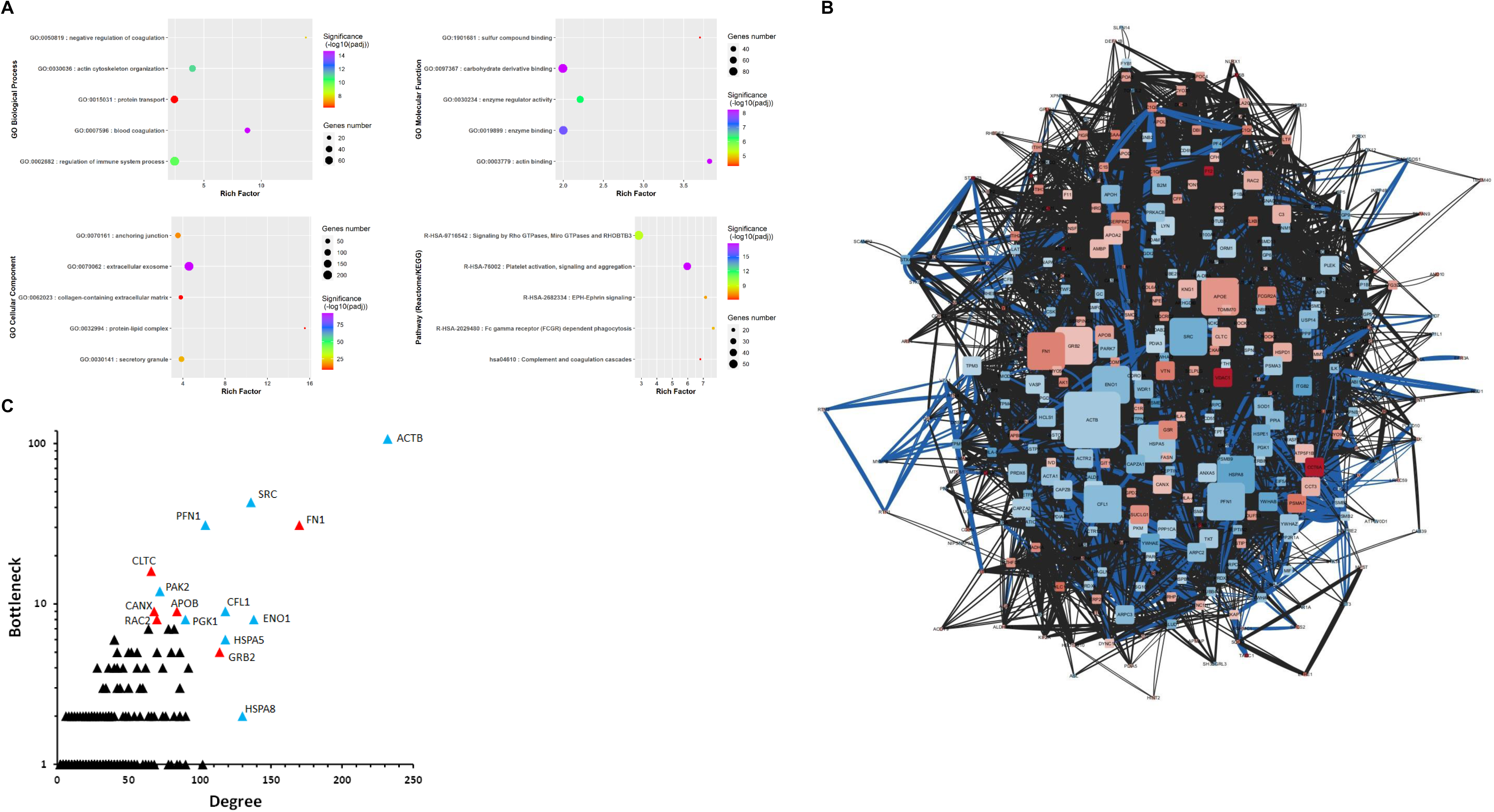
Functional analysis and network building from 369 differentially detected proteins in EVs purified from TDV compared to PV. (A) Top-5 of more enriched terms of Gene Ontology Biological Process (BP), Molecular Function (MF) and Cellular Component (CC) and associated to pathway database (Reactome and KEGG). Enrichment analysis was performed on Database for Annotation, Visualization and Integrated Discovery (DAVID) online tool (https://david.ncifcrf.gov/) with cluster analysis of enriched terms performing to recover redundant terms. The selected term corresponds to the one with the most significant p-adjusted within each top-5 enrichment cluster. (B) Protein-protein interaction between 369 proteins was retrieved from Search Tool for the Retrieval of Interacting Genes/Proteins (STRING) database and used to build interaction network with Cytoscape environment. Among 369 proteins identified by LC-MS/MS, 361 matched onto STRING database. Densely connected network was obtained with 5492 edges (interactions) between 342 nodes (proteins). Thickness of edges correspond to confidence level, a score between 0 to 1, calculated by STRING from different sources of information to identify protein-protein interaction. Blue colors of edges highlight physical interaction, supported by experimental data from interactome database. Color gradient from blue to red correspond to fold change of down- and up-regulated proteins of EVs from TDV *vs* EVs from PV, respectively. Size of nodes is proportional to degree of node, *i.e* the number of interactant of node. (C) Topological parameters analysis on network building from the functional association between proteins deregulated in EVs from TDV *vs.* EVs from PV were calculated by Cytohubba, a Cytoscape plug-in that permit to predict important nodes for the network, and by extension, important protein in biological context. Scatter plot displaying the correlation between two topological parameters, node degree and bottleneck (a centrality parameter based on shortest paths). Official symbol and level of deregulation (blue for downregulation and red for upregulation) were indicated for important node, *i.e* displaying high values for degree and bottleneck.

The significance of these protein clusters was confirmed in the network showing protein-protein interaction (Figure 7B). Densely connected network was obtained with 5492 edges (interactions) between 342 nodes (proteins). The density of this network suggested that all proteins are closely and functionally linked and thus belonged or are involved in similar functions or pathways supporting previous observations achieved with GO.

The top 15 hub proteins, including 6 upregulated proteins (FN1, CLTC, CANX, APOB, RAC2, GRB2) and 9 downregulated proteins (ACTB, SRC, PFN1, PAK2, CFL1, PGK1, ENO1, HSPA5, HSPA8) were identified according to the highest degrees of connectivity using cytoHubba the plug-in in Cytoscape (Figure 7C). Among upregulated proteins, a single (APOB) protein directly linked to cholesterol transport, was previously demonstrated to be overexpressed in EVs from TDV whereas it was underexpressed in the tumor and, rather linked to good prognosis. All other proteins were associated with a poor prognosis in lung cancer.

## DISCUSSION

The development of liquid biopsy constitutes a crucial breakthrough for identifying potential tumor biomarkers and thus monitoring patients during treatment in order to limit recurrences or prevent therapeutic escape. In this context, EVs and their content afford a putative and promising source of circulating biomarkers. However, as already demonstrated in previous study, the cargo of EVs might be variable according to the sampling site [18]. Indeed, previous work has demonstrated an original miRNA signature in EVs derived from TDV. The most expressed miRNA of this signature, miR-203a-3p, was significantly higher in EVs derived from TDV compared to peripheral blood EVs [18]. Nevertheless, to date, no study has demonstrated any differences between proteins isolated in central EVs and those derived from PV. Navarro *et al*. have described that the size of EVs could be modified according to the site of sampling, from pulmonary or peripheral veins (mean mode size 111.9 vs 118.2) but not the concentration (3.85 10^9^ particles/mL) [16]. In this study, we decided to investigate EVs size and concentration. The concentration of EVs was significantly higher in TDV plasma samples compared as to those in peripheral samples (17.0 10^9^ vs 4.4 10^9^ particles/mL respectively). On the contrary, TDV from EVs were significantly smaller than EVs from PV (mean mode size: 74.7 vs 97.2 nm).

Strikingly, we showed that pulmonary vein EVs concentration was 4-fold higher than the Navarro’s series, whereas peripheral EVs concentration was similar. Moreover, both TDV and PV EVs were smaller in our series. These differences were neither explained by the EVs isolation and characterization methods (ultracentrifugation and then NTA using Nanosight NS-300), nor the pathological tumor stage (stage I: 45% vs 49%, stage II-III: 55% versus 51% in our series vs Navarro *et al.* series respectively). However, we studied only adenocarcinomas whereas previous series included around 60% of adenocarcinomas, 25% of squamous cell carcinomas and 15% of other lung cancer types, which could influence characteristics of EVs in the tumor-draining vein [18].

TDV samples were enriched with smaller EVs, with approximately 3 times more EVs with a mean size <100 nm than those with a mean size >100 nm. This difference was not observed in PV samples. Navarro *et al.* described only a higher proportion of very small exosomes (<50nm of size) in pulmonary vein *vs* peripheral vein [16].

Concentration of EVs in TDV was not correlated with the size of the tumor in our cohort whereas previous work revealed a significant increase of TDV EVs concentration according to the tumor size. As already demonstrated, a smaller size of EVs in pulmonary vein was significantly associated with relapse and survival of patients, even in stage-I cancers [16]. Thus, the size threshold to assess high-risk of recurrence patients was inferior to 112 nm. In our study, mean mode size of EVs purified from TDV was 75 nm, but did not impact the risk of relapse. However, the very low number of recurrence cases (n=4) in our study was not sufficient to establish a link between EVs size and aggressiveness of the tumor.

In a second step, the comparison of proteomic profiling of EVs derived from TDV and PV plasma samples revealed that 10 proteins were overexpressed in EVs from TDV compared to those from PV. Nevertheless, only one protein (SRPRB) was overexpressed in the TDV-derived EVs compared to both PV-derived EVs and tumor compared to paired non-tumoral tissues. SRPRB correspond to a signal recognition particle receptor β that has already been detected in endothelial cell-derived EVs. The transfer of SRPRB in EVs alters vascular smooth muscle cell phenotype [34]. Since this protein is involved in sorting secretory and membrane proteins to the endoplasmic reticulum membranes, its upregulation in EVs from TDV and tumor cells suggests an increase of membrane protein synthesis and excretion [35]. SRPRB was expressed in most of the half of lung tumors and is significantly associated with tumor progression and bad prognosis. Indeed, its function was clearly linked to regulation of cell proliferation, apoptosis, tumorigenesis and metastasis [36]. In this context, the release of SRPRB in early stages tumor-derived EVs could contribute to the dissemination of tumor biomarkers to control the tumor microenvironment. Consequently, Among the 10 most specific proteins of EVs from TDV, 9 were associated with diagnosis and prognosis of lung cancer (CCT6A, MUC5B, VDAC1, ATP1A1, F12) and other cancer subtypes (UNC45A, SSC5D, IGHV5-51). These proteins play a major role in tumor progression and are directly associated with poor prognosis in NSCLC. Thus, CCT6A plays a key role in lung adenocarcinoma progression promoting the stabilization of STAT1 and inducing metabolic reprogramming [37]. Similarly, high expression of MUC5B mRNA was significantly associated with poor OS and progression free survival in patients with lung adenocarcinoma [38, 39] EVs also appear to carry mitochondrial voltage-dependent anion channel (VDAC) whose function lies at the crossroads between metabolic and survival pathways, and thus contribute to oncogenesis and metastasis [40]. Since EV from patients with cancer express increased amounts of polyphosphate (PolyP) compared with healthy individuals’ polyphosphate and that PolyP-expressing EVs binds the coagulation factor XII (F12), its upregulation in EVs from TDV suggest that these EVs are derived from tumor cells [41]. In contrast to previous observations, we found that 12 proteins were overexpressed in EVs from TDV whereas they were underexpressed in tumor (i.e. specific to normal lung tissue): AHNAK, ITIH2, APOB, SERPING1, APOA2, COL6A1, MCEMP1, PCYOX1, PGLYRP2, HLA-E, CSRP1, APOC1. Strinkingly, among them, five were known to be down-expressed in lung cancer tissue or presented a tumor suppressive function (AHNAK, APOB, ITIH2, MCEMP1, COL6A1). We hypothesize that lung tumor cells were able to release these tumor suppressive proteins, as demonstrated with the gastrokine 1. This protein is a stomach-specific tumor suppressor that is secreted into extracellular space as an exosomal cargo protein and inhibits gastric oncogenesis [42].

To the best of our knowledge, this is the first comprehensive and comparative proteomic analysis of circulating EVs derived from TDV plasma samples and those from PV of patients with locally advanced NSCLC. We identified herein putative proteomic signatures, that might represent novel circulating biomarkers for NSCLC prognosis and predicting recurrences at early stages.

