## Supplementary data for "New proteomic signature in circulating extracellular vesicles from tumor-draining vein of lung adenocarcinomas patients"

### Supplementary figure legends

**Supplementary figure 1.** Venn diagram of 1410 proteins identified in at least 70% of the EVs samples (from Tumor-draining vein and Peripheral vein) compared with 8452 proteins annotated in the Vesiclepedia (https://www.microvesicles.org/) and 6517 human proteins annotated in Exocarta (http://exocarta.org/) databases.

**Supplementary figure 2: Prognosis potential of protein detection in EVs purified from TDV or PV plasma samples during recurrence.**

Differential analysis was conducted from EVs isolated from Tumor-draining vein (TDV) (A) and from EVs isolated from Peripheral vein (PV) (B) according to recurrence. Deregulated proteins were selected using thresholds $|(\log_2 FC| \geq 1$ and p value $\leq 0.05$, due to loss of statistical power when comparing groups with unbalanced numbers of samples: 4 recurrent versus 15 no recurrent tumor for TDV (A) and 3 recurrent versus 11 no recurrent tumor for PV (B). Hierarchical clustering analysis was conducting from 80 differentially detected protein in EVs from TDV of recurrent tumor as compared to no recurrent tumor (A) and from 92 differentially detected protein in EVs from PV of recurrent tumor as compared to no recurrent tumor (B). Protein intensities were log2 transformed and median-centered and are displayed as color gradient from blue to red, reflecting low to high protein level in EVs, respectively. Proteins (rows) and tissue samples (columns) are hierarchically clustered using Pearson correlation distance and average linkage method. Samples characteristics were indicated at the top, with gender distribution (male: blue, female: pink), stage (early stage composed IA, IB and IIA pTNM staging: yellow, intermediate/late stage represented by IIB and IIIA pTNM staging: violet) and recurrence (no: white, yes: black).

**Supplementary figure 1.**

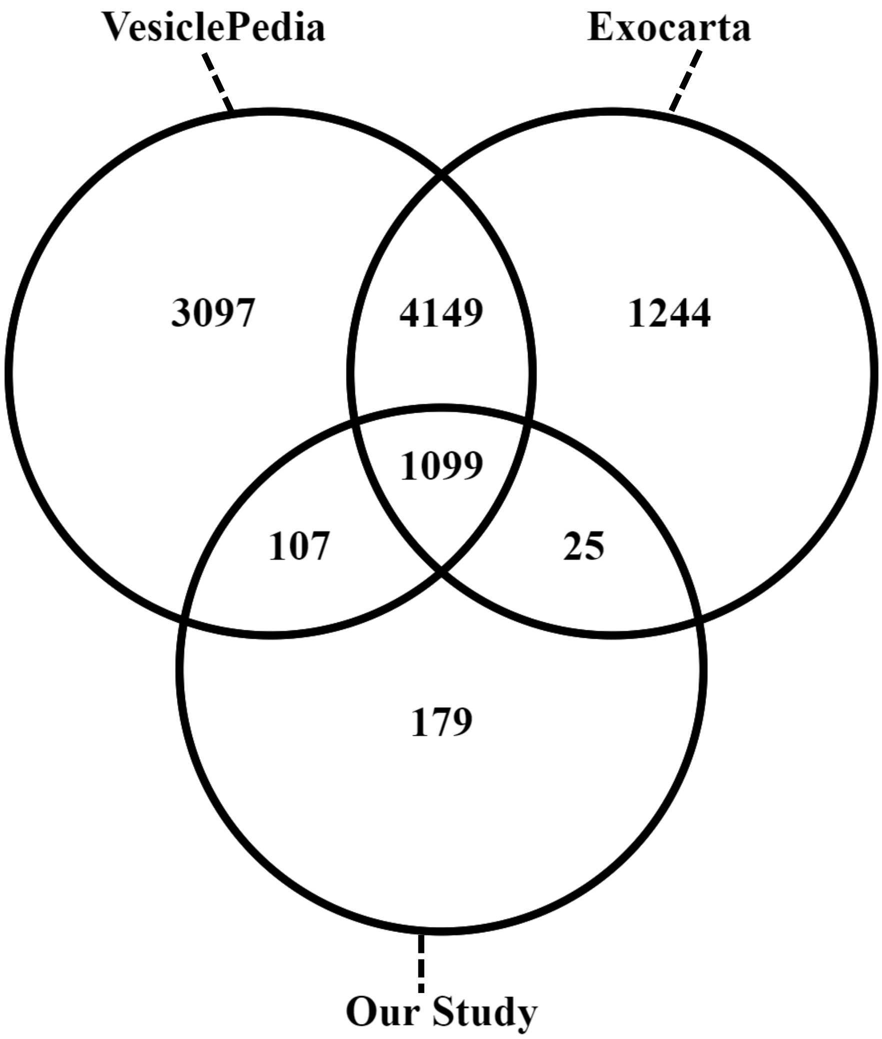

**Supplementary figure 2.**

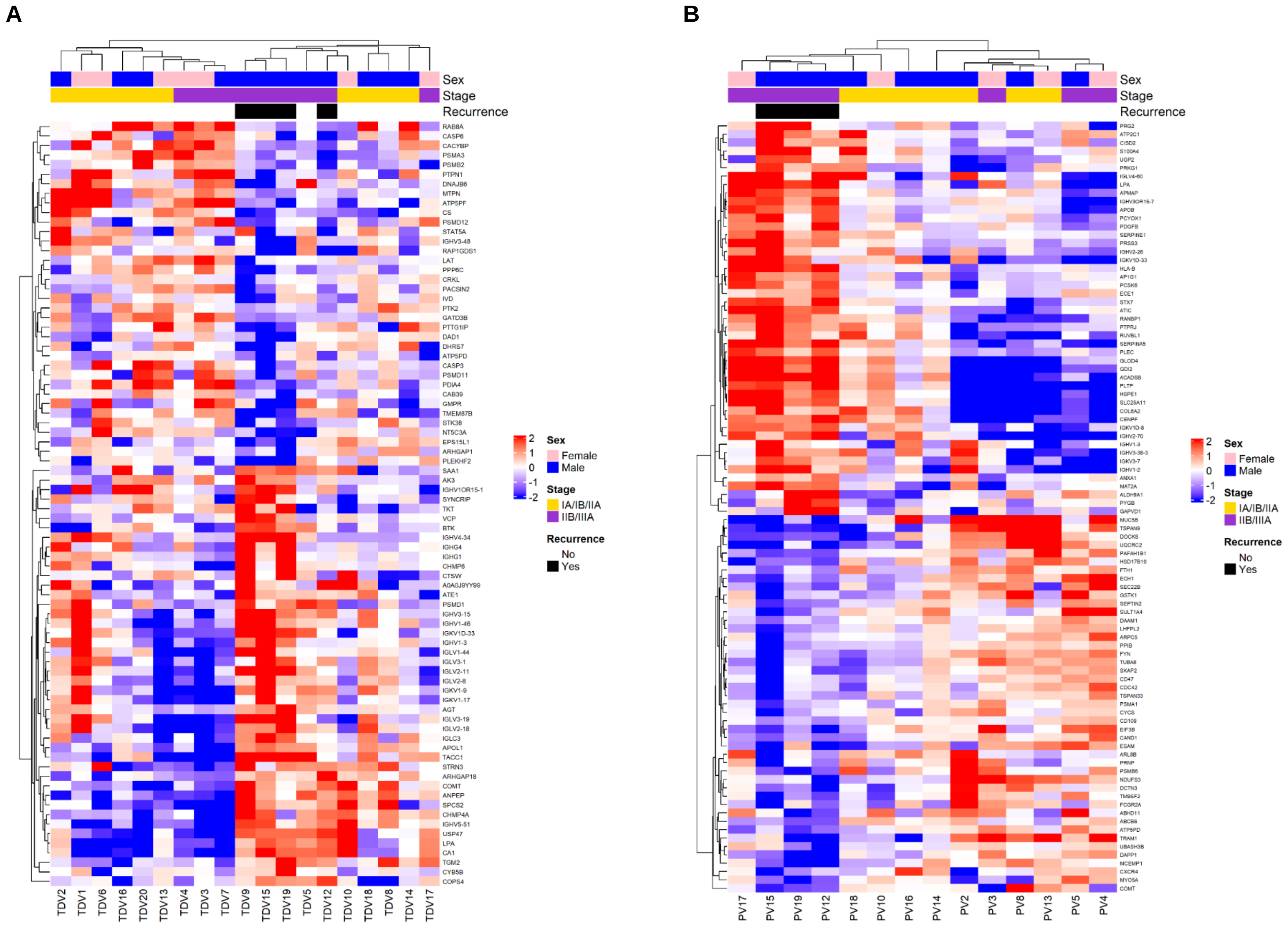

**Supplementary Table 1.** Deregulated proteins in lung adenocarcinoma as compared to non-tumoral tissues identified by mass spectrometry, according to the following thresholds: |log2 fold change| ≥ 1 and adjusted p value (Benjamini-Hochberg multiple testing correction) ≤ 0.05.

| Official Gene Symbol | Uniprot ID | $log_2$ Fold Change (T vs NT) | adjusted p value |
|---|---|---|---|
| PYCR1 | P32322 | 4,04 | 9,43E-09 |
| GOLM1 | Q8NBJ4 | 2,97 | 1,37E-07 |
| TUBB3 | Q13509 | 2,42 | 3,38E-05 |
| S100B | P04271 | 2,38 | 4,35E-05 |
| TMEM165 | Q9HC07 | 2,3 | 3,49E-06 |
| CYP1B1 | Q16678 | 2,29 | 0,001765407 |
| OCIAD2 | Q56VL3 | 2,29 | 1,37E-05 |
| CPD | O75976 | 2,26 | 0,000127022 |
| UGT1A6 | P19224 | 2,25 | 0,006993255 |
| PES1 | O00541 | 2,2 | 2,97E-08 |
| MZB1 | Q8WU39 | 2,18 | 0,002047982 |
| RBM47 | A0AV96 | 2,17 | 1,81E-08 |
| FCGBP | Q9Y6R7 | 2,12 | 0,000130139 |
| MCM3 | P25205 | 2,11 | 3,25E-05 |
| PDLIM4 | P50479 | 1,93 | 4,03E-06 |
| CTHRC1 | Q96CG8 | 1,92 | 0,000118446 |
| LAD1 | O00515 | 1,91 | 4,11E-05 |
| DCTPP1 | Q9H773 | 1,86 | 7,66E-06 |
| TUBA1C | Q9BQE3 | 1,79 | 6,94E-06 |
| NUP85 | Q9BW27 | 1,78 | 2,54E-05 |
| FKBP11 | Q9NYL4 | 1,77 | 2,68E-05 |
| GALNT2 | Q10471 | 1,77 | 6,93E-06 |
| PPP1R14B | Q96C90 | 1,75 | 9,43E-07 |
| GFPT1 | Q06210 | 1,74 | 2,10E-06 |
| CASP6 | P55212 | 1,73 | 2,03E-06 |
| NEU1 | Q99519 | 1,73 | 5,33E-05 |
| CRABP2 | P29373 | 1,72 | 0,000138023 |
| PC | P11498 | 1,72 | 3,95E-06 |
| NQO1 | P15559 | 1,68 | 0,005969867 |
| BCS1L | Q9Y276 | 1,64 | 5,94E-05 |
| GALNT3 | Q14435 | 1,64 | 6,83E-05 |
| PTRH2 | Q9Y3E5 | 1,63 | 5,86E-05 |
| SDF2L1 | Q9HCN8 | 1,59 | 5,07E-05 |
| EIF5B | O60841 | 1,58 | 9,92E-05 |
| SPATS2L | Q9NUQ6 | 1,57 | 0,000892923 |
| ADI1 | Q9BV57 | 1,55 | 4,91E-05 |
| AKR1C1 | Q04828 | 1,55 | 0,010839734 |
| FAP | Q12884 | 1,55 | 0,003825763 |
| PAK1 | Q13153 | 1,55 | 6,45E-05 |
| SLC35B2 | Q8TB61 | 1,55 | 1,38E-05 |

| | | | |
|---|---|---|---|
| RCN3 | Q96D15 | 1,54 | 0,001170267 |
| NLN | Q9BYT8 | 1,5 | 7,31E-06 |
| FEN1 | P39748 | 1,47 | 0,001196246 |
| UBQLN1 | Q9UMX0 | 1,47 | 0,000891738 |
| ALDH1A3 | P47895 | 1,46 | 0,010225178 |
| CRTAP | O75718 | 1,46 | 0,001369952 |
| EIF2B5 | Q13144 | 1,46 | 2,31E-06 |
| FAM3C | Q92520 | 1,46 | 9,68E-05 |
| SERPINH1 | P50454 | 1,46 | 0,000747569 |
| FAU | P62861 | 1,45 | 0,000412122 |
| AGR2 | O95994 | 1,44 | 0,002598472 |
| UCHL1 | P09936 | 1,43 | 0,00290664 |
| UGDH | O60701 | 1,42 | 0,000751072 |
| CHORDC1 | Q9UHD1 | 1,41 | 0,000684891 |
| ALDH18A1 | P54886 | 1,4 | 9,54E-07 |
| SLC25A10 | Q9UBX3 | 1,4 | 3,09E-06 |
| BCCIP | Q9P287 | 1,38 | 4,11E-05 |
| RPL19 | P84098 | 1,37 | 0,001760542 |
| SEC14L2 | O76054 | 1,37 | 0,001003813 |
| MAN1B1 | Q9UKM7 | 1,36 | 3,67E-05 |
| TNC | P24821 | 1,36 | 0,012732216 |
| RPL24 | P83731 | 1,35 | 2,63E-05 |
| SLC3A2 | P08195 | 1,34 | 0,003663465 |
| SMAD4 | Q13485 | 1,34 | 0,00051138 |
| CD38 | P28907 | 1,32 | 0,048520318 |
| ISG15 | P05161 | 1,32 | 0,000655838 |
| MCM6 | Q14566 | 1,31 | 0,000149968 |
| STEAP3 | Q658P3 | 1,31 | 1,44E-06 |
| CLPTM1 | O96005 | 1,3 | 0,00047677 |
| FKBP10 | Q96AY3 | 1,3 | 0,013941166 |
| NT5C3A | Q9H0P0 | 1,3 | 0,000425794 |
| RPL23 | P62829 | 1,3 | 0,003171311 |
| COG3 | Q96JB2 | 1,29 | 9,43E-07 |
| FUT8 | Q9BYC5 | 1,29 | 8,13E-06 |
| HDHD3 | Q9BSH5 | 1,29 | 0,000110565 |
| TPMT | P51580 | 1,29 | 0,00058922 |
| SLC16A3 | O15427 | 1,28 | 6,80E-05 |
| AP1M2 | Q9Y6Q5 | 1,25 | 0,00014121 |
| RPL27A | P46776 | 1,24 | 4,98E-06 |
| RSL1D1 | O76021 | 1,24 | 5,24E-05 |
| AGA | P20933 | 1,22 | 0,007645818 |
| CDV3 | Q9UKY7 | 1,22 | 0,000369585 |
| IL4I1 | Q96RQ9 | 1,22 | 0,005753725 |
| NSUN2 | Q08J23 | 1,22 | 1,18E-05 |
| PDXDC1 | Q6P996 | 1,22 | 0,000127459 |

| | | | |
|---|---|---|---|
| SNAP29 | O95721 | 1,22 | 0,00313284 |
| SRPRB | Q9Y5M8 | 1,22 | 5,55E-06 |
| GLB1 | P16278 | 1,21 | 0,002543473 |
| PFKP | Q01813 | 1,21 | 8,50E-05 |
| SPINT2 | O43291 | 1,21 | 0,000841601 |
| TMCO1 | Q9UM00 | 1,21 | 0,000343997 |
| FAHD1 | Q6P587 | 1,2 | 0,003246611 |
| CIT | O14578 | 1,18 | 0,017651986 |
| GALE | Q14376 | 1,18 | 1,23E-05 |
| ERO1A | Q96HE7 | 1,17 | 4,15E-05 |
| NAXE | Q8NCW5 | 1,17 | 2,37E-06 |
| POSTN | Q15063 | 1,17 | 0,003940669 |
| SHMT2 | P34897 | 1,17 | 2,50E-05 |
| BPNT2 | Q9NX62 | 1,16 | 0,000378755 |
| BZW2 | Q9Y6E2 | 1,16 | 0,000415456 |
| DKC1 | O60832 | 1,16 | 0,002240309 |
| NAMPT | P43490 | 1,16 | 0,00129936 |
| PLOD1 | Q02809 | 1,16 | 0,006508024 |
| PSME3 | P61289 | 1,16 | 4,21E-06 |
| SURF4 | O15260 | 1,16 | 0,000327947 |
| B4GALT1 | P15291 | 1,15 | 1,40E-06 |
| EIF3K | Q9UBQ5 | 1,15 | 0,008819761 |
| HID1 | Q8IV36 | 1,15 | 4,17E-05 |
| NIPSNAP1 | Q9BPW8 | 1,15 | 0,00108148 |
| RPL31 | P62899 | 1,14 | 2,58E-05 |
| ADGRE5 | P48960 | 1,13 | 0,002909721 |
| ARFGAP1 | Q8N6T3 | 1,13 | 0,000179855 |
| DDX21 | Q9NR30 | 1,13 | 0,00029923 |
| OAT | P04181 | 1,12 | 0,006969032 |
| ARFGEF1 | Q9Y6D6 | 1,11 | 0,000148136 |
| FTSJ3 | Q8IY81 | 1,09 | 0,000792568 |
| RRP12 | Q5JTH9 | 1,09 | 0,00102395 |
| PUS1 | Q9Y606 | 1,08 | 0,000546641 |
| SERPINB5 | P36952 | 1,08 | 0,019109927 |
| TMED3 | Q9Y3Q3 | 1,08 | 0,000136694 |
| TMTC3 | Q6ZXV5 | 1,08 | 0,000115609 |
| IDE | P14735 | 1,07 | 0,000527262 |
| MYBBP1A | Q9BQG0 | 1,07 | 0,000207434 |
| SGPL1 | O95470 | 1,07 | 5,77E-05 |
| TMEM87B | Q96K49 | 1,07 | 0,001928988 |
| AMACR | Q9UHK6 | 1,06 | 0,002010974 |
| NOL3 | O60936 | 1,06 | 5,88E-05 |
| S100P | P25815 | 1,06 | 0,031170776 |
| SEC23IP | Q9Y6Y8 | 1,06 | 0,010397674 |
| HM13 | Q8TCT9 | 1,05 | 0,00011223 |

| | | | |
|---|---|---|---|
| DSG2 | Q14126 | 1,04 | 0,02662929 |
| HSP90B2P | Q58FF3 | 1,04 | 0,000138382 |
| MAN2A1 | Q16706 | 1,04 | 0,007822015 |
| RAB9A | P51151 | 1,04 | 0,018227802 |
| RPL7A | P62424 | 1,04 | 2,34E-06 |
| ARFIP1 | P53367 | 1,03 | 0,00290664 |
| FAM114A2 | Q9NRY5 | 1,03 | 0,000724574 |
| RAB25 | P57735 | 1,03 | 0,000111636 |
| SLC16A4 | O15374 | 1,03 | 0,000461579 |
| TMEM214 | Q6NUQ4 | 1,03 | 0,004631536 |
| LSR | Q86X29 | 1,02 | 0,000655087 |
| CRELD2 | Q6UXH1 | 1,01 | 0,00051181 |
| DUSP23 | Q9BVJ7 | 1,01 | 0,000232456 |
| FUBP3 | Q96I24 | 1,01 | 0,001580465 |
| GLA | P06280 | 1,01 | 0,037578152 |
| NAE1 | Q13564 | 1,01 | 0,000769851 |
| PPIC | P45877 | 1,01 | 0,000197634 |
| RPS17 | P08708 | 1,01 | 8,78E-06 |
| ABCA3 | Q99758 | -1,01 | 0,000620952 |
| FGA | P02671 | -1,01 | 0,018266214 |
| PGLYRP2 | Q96PD5 | -1,01 | 0,000236336 |
| PTPRG | P23470 | -1,01 | 2,54E-05 |
| VCL | P18206 | -1,01 | 5,98E-06 |
| ATP5PD | O75947 | -1,02 | 3,49E-06 |
| CYP27A1 | Q02318 | -1,02 | 0,001352079 |
| HCFC1 | P51610 | -1,02 | 0,000150224 |
| MTARC2 | Q969Z3 | -1,02 | 0,000200261 |
| RAC1 | P63000 | -1,02 | 2,55E-06 |
| ZYX | Q15942 | -1,02 | 3,48E-05 |
| H3-7 | Q5TEC6 | -1,03 | 0,001764671 |
| MXRA7 | P84157 | -1,03 | 9,92E-05 |
| CD82 | P27701 | -1,04 | 0,012638938 |
| GNAQ | P50148 | -1,04 | 0,000347462 |
| HNRNPH3 | P31942 | -1,04 | 0,007168145 |
| SEPTIN11 | Q9NVA2 | -1,04 | 6,89E-07 |
| APOA4 | P06727 | -1,05 | 0,003670017 |
| GPX3 | P22352 | -1,05 | 0,000574265 |
| ITGA6 | P23229 | -1,05 | 0,009391907 |
| NEK7 | Q8TDX7 | -1,05 | 0,001765407 |
| PCYOX1 | Q9UHG3 | -1,05 | 3,49E-06 |
| BTN3A2 | P78410 | -1,06 | 6,78E-05 |
| FERMT2 | Q96AC1 | -1,06 | 0,003765036 |
| HLA-E | P13747 | -1,06 | 0,000412585 |
| LLGL2 | Q6P1M3 | -1,06 | 0,000140503 |
| TF | P02787 | -1,06 | 0,00392286 |

| | | | |
|---|---|---|---|
| CD59 | P13987 | -1,07 | 2,67E-05 |
| FCN3 | O75636 | -1,07 | 0,000763763 |
| ADD1 | P35611 | -1,08 | 3,32E-05 |
| BCAM | P50895 | -1,08 | 1,41E-05 |
| CHMP1B | Q7LBR1 | -1,08 | 0,001428795 |
| FLNC | Q14315 | -1,08 | 0,014078388 |
| KCTD12 | Q96CX2 | -1,08 | 1,42E-06 |
| SVIL | O95425 | -1,08 | 0,0304392 |
| TRIP10 | Q15642 | -1,08 | 8,11E-05 |
| VWF | P04275 | -1,08 | 6,36E-05 |
| AGT | P01019 | -1,09 | 0,024154148 |
| CSRP1 | P21291 | -1,09 | 0,001097241 |
| DDAH2 | O95865 | -1,09 | 9,45E-06 |
| SH3D19 | Q5HYK7 | -1,09 | 6,52E-06 |
| TMEM119 | Q4V9L6 | -1,09 | 9,68E-07 |
| ALDH1A1 | P00352 | -1,1 | 0,000127022 |
| DNAJB4 | Q9UDY4 | -1,1 | 0,011796993 |
| POLD1 | P28340 | -1,1 | 8,54E-06 |
| RAB8B | Q92930 | -1,1 | 2,55E-06 |
| SEPTIN10 | Q9P0V9 | -1,1 | 2,39E-05 |
| CALCRL | Q16602 | -1,11 | 0,015007641 |
| IFIT3 | O14879 | -1,11 | 0,007814608 |
| CADM1 | Q9BY67 | -1,12 | 0,006919724 |
| ISYNA1 | Q9NPH2 | -1,12 | 0,004402734 |
| SARG | Q9BW04 | -1,12 | 0,001908428 |
| SNTB2 | Q13425 | -1,12 | 0,006014462 |
| CRIP2 | P52943 | -1,13 | 0,000819457 |
| CYB5A | P00167 | -1,13 | 6,33E-05 |
| ITIH2 | P19823 | -1,13 | 0,000686435 |
| MPP1 | Q00013 | -1,13 | 0,000269519 |
| NEBL | O76041 | -1,13 | 3,03E-05 |
| SELENOT | P62341 | -1,13 | 0,000583421 |
| VIM | P08670 | -1,13 | 5,68E-05 |
| APCS | P02743 | -1,14 | 0,00215016 |
| MYL9 | P24844 | -1,14 | 0,000945769 |
| PALMD | Q9NP74 | -1,14 | 0,000413097 |
| APOB | P04114 | -1,15 | 0,000519221 |
| CD5L | O43866 | -1,15 | 0,033171861 |
| H1-2 | P16403 | -1,15 | 1,33E-05 |
| H1-4 | P10412 | -1,15 | 1,33E-05 |
| LSAMP | Q13449 | -1,15 | 8,33E-06 |
| RALA | P11233 | -1,15 | 5,33E-05 |
| AK4 | P27144 | -1,17 | 0,002340981 |
| AP3B1 | O00203 | -1,17 | 3,15E-07 |
| CRMP1 | Q14194 | -1,17 | 0,000327947 |

| | | | |
|---|---|---|---|
| FGB | P02675 | -1,17 | 0,009336211 |
| KRT20 | P35900 | -1,17 | 0,029451921 |
| PDLIM3 | Q53GG5 | -1,17 | 0,006926868 |
| APOC1 | P02654 | -1,18 | 0,007960582 |
| EPB41L3 | Q9Y2J2 | -1,18 | 0,000346648 |
| CD9 | P21926 | -1,19 | 2,22E-05 |
| GLIPR2 | Q9H4G4 | -1,19 | 6,59E-06 |
| H2AC20 | Q16777 | -1,19 | 6,95E-05 |
| SERPING1 | P05155 | -1,19 | 0,001499669 |
| ACP5 | P13686 | -1,2 | 0,022396581 |
| AMPD3 | Q01432 | -1,2 | 0,001105961 |
| CD55 | P08174 | -1,2 | 0,000577279 |
| PLVAP | Q9BX97 | -1,2 | 0,000421827 |
| CACNA2D2 | Q9NY47 | -1,21 | 0,009992923 |
| CYGB | Q8WWM9 | -1,21 | 0,000207598 |
| DOCK4 | Q8N1I0 | -1,21 | 1,52E-05 |
| FXYD1 | O00168 | -1,21 | 0,001584332 |
| MYO1C | O00159 | -1,21 | 5,32E-07 |
| SORBS2 | O94875 | -1,21 | 0,031504829 |
| SULT1A4 | P0DMN0 | -1,21 | 0,007535331 |
| COL6A6 | A6NMZ7 | -1,22 | 0,003399807 |
| HPGD | P15428 | -1,22 | 0,032004054 |
| MCEMP1 | Q8IX19 | -1,22 | 0,008819761 |
| RDX | P35241 | -1,22 | 7,61E-07 |
| CNN1 | P51911 | -1,23 | 0,033396288 |
| RHD | Q02161 | -1,23 | 0,001124664 |
| PRTN3 | P24158 | -1,24 | 0,019663556 |
| SNCA | P37840 | -1,24 | 2,74E-05 |
| DPYSL2 | Q16555 | -1,25 | 2,73E-07 |
| EHD4 | Q9H223 | -1,25 | 5,77E-07 |
| ERN1 | O75460 | -1,25 | 0,002653604 |
| HP | P00738 | -1,25 | 0,033396288 |
| MYLK | Q15746 | -1,25 | 0,000197536 |
| PLPP3 | O14495 | -1,25 | 6,87E-06 |
| GTF2IRD2 | Q86UP8 | -1,26 | 0,002632009 |
| SULT1A1 | P50225 | -1,26 | 0,00151811 |
| TPSB2 | P20231 | -1,26 | 0,002970004 |
| LUM | P51884 | -1,27 | 0,000237946 |
| OCLN | Q16625 | -1,27 | 0,001002152 |
| PRKG1 | Q13976 | -1,27 | 0,000184112 |
| RECK | O95980 | -1,27 | 0,000141221 |
| ANXA2 | P07355 | -1,28 | 7,82E-07 |
| IRF4 | Q15306 | -1,28 | 0,002593333 |
| MACF1 | Q9UPN3 | -1,28 | 1,37E-07 |
| RALB | P11234 | -1,28 | 0,001810375 |

| | | | |
|---|---|---|---|
| SND1 | Q7KZF4 | -1,28 | 0,001680803 |
| TJP1 | Q07157 | -1,28 | 5,20E-07 |
| USO1 | O60763 | -1,28 | 0,000401882 |
| DCN | P07585 | -1,29 | 0,001337651 |
| STOM | P27105 | -1,29 | 2,39E-05 |
| TRMT61A | Q96FX7 | -1,29 | 0,003407076 |
| ANXA9 | O76027 | -1,3 | 1,72E-06 |
| GUCY1B1 | Q02153 | -1,3 | 7,03E-06 |
| RTN3 | O95197 | -1,3 | 0,000213943 |
| ANXA13 | P27216 | -1,31 | 4,00E-07 |
| LAMP3 | Q9UQV4 | -1,31 | 0,000139364 |
| LMOD1 | P29536 | -1,31 | 0,002744414 |
| SAMM50 | Q9Y512 | -1,31 | 0,001039787 |
| WFS1 | O76024 | -1,31 | 6,89E-07 |
| FHOD1 | Q9Y613 | -1,32 | 3,96E-05 |
| ITGA3 | P26006 | -1,32 | 1,78E-05 |
| KRT4 | P19013 | -1,32 | 0,000718514 |
| S100A4 | P26447 | -1,32 | 1,64E-05 |
| SPON1 | Q9HCB6 | -1,32 | 0,000197536 |
| FABP5 | Q01469 | -1,33 | 8,03E-05 |
| MYADM | Q96S97 | -1,33 | 1,39E-05 |
| PARVA | Q9NVD7 | -1,33 | 0,000138382 |
| SLC30A9 | Q6PML9 | -1,33 | 0,000168151 |
| ASPM | Q8IZT6 | -1,34 | 0,001731264 |
| MYH10 | P35580 | -1,34 | 4,35E-05 |
| S100A12 | P80511 | -1,34 | 0,008240675 |
| SERPIND1 | P05546 | -1,34 | 0,001045397 |
| ITIH4 | Q14624 | -1,35 | 0,002625667 |
| PXMP4 | Q9Y6I8 | -1,35 | 0,000195913 |
| SERPINA6 | P08185 | -1,35 | 0,006822668 |
| AFM | P43652 | -1,37 | 0,002527921 |
| CD81 | P60033 | -1,37 | 1,18E-05 |
| DDT | P30046 | -1,37 | 1,72E-06 |
| PSPC1 | Q8WXF1 | -1,37 | 0,000332987 |
| S100A10 | P60903 | -1,37 | 9,43E-07 |
| AOX1 | Q06278 | -1,38 | 0,0012715 |
| COL6A1 | P12109 | -1,38 | 0,011763271 |
| NEXN | Q0ZGT2 | -1,38 | 0,001931642 |
| LMCD1 | Q9NZU5 | -1,39 | 0,000127459 |
| SPTAN1 | Q13813 | -1,39 | 1,80E-06 |
| ABLIM1 | O14639 | -1,4 | 0,000480349 |
| CFD | P00746 | -1,4 | 0,006205252 |
| HCFC2 | Q9Y5Z7 | -1,4 | 0,000432369 |
| SLC44A2 | Q8IWA5 | -1,4 | 2,61E-05 |
| AHNAK | Q09666 | -1,41 | 2,73E-07 |

| | | | |
|---|---|---|---|
| TPM2 | P07951 | -1,41 | 0,007234638 |
| APOA1 | P02647 | -1,42 | 0,001249824 |
| AQP1 | P29972 | -1,42 | 0,000819428 |
| MAB21L4 | Q08AI8 | -1,42 | 0,000138382 |
| ANO6 | Q4KMQ2 | -1,43 | 1,18E-06 |
| EPB41 | P11171 | -1,43 | 2,85E-05 |
| STXBP1 | P61764 | -1,43 | 0,000281579 |
| APOA2 | P02652 | -1,44 | 0,002211648 |
| ARMC1 | Q9NVT9 | -1,45 | 1,28E-05 |
| ADD3 | Q9UEY8 | -1,47 | 0,000197634 |
| DES | P17661 | -1,47 | 0,00079089 |
| GNAI1 | P63096 | -1,47 | 8,97E-05 |
| PRDX2 | P32119 | -1,47 | 0,002017188 |
| BST2 | Q10589 | -1,48 | 0,000211672 |
| PPBP | P02775 | -1,48 | 0,001760598 |
| A2M | P01023 | -1,49 | 4,11E-05 |
| CA3 | P07451 | -1,49 | 6,73E-05 |
| PKP3 | Q9Y446 | -1,49 | 0,002803928 |
| PRELP | P51888 | -1,49 | 0,000184281 |
| TSPAN8 | P19075 | -1,49 | 0,005119711 |
| F9 | P00740 | -1,5 | 1,45E-06 |
| HSD17B6 | O14756 | -1,5 | 0,007960582 |
| RRAS2 | P62070 | -1,5 | 2,30E-07 |
| DNAJB6 | O75190 | -1,51 | 0,000107997 |
| FBLN5 | Q9UBX5 | -1,51 | 0,000206618 |
| EHD3 | Q9NZN3 | -1,52 | 1,48E-05 |
| LMO7 | Q8WWI1 | -1,52 | 5,32E-07 |
| CKB | P12277 | -1,53 | 9,52E-05 |
| KANK2 | Q63ZY3 | -1,53 | 6,70E-06 |
| COL6A2 | P12110 | -1,54 | 0,007811501 |
| MAOB | P27338 | -1,54 | 1,72E-06 |
| CD151 | P48509 | -1,55 | 7,47E-05 |
| RRAS | P10301 | -1,57 | 3,49E-06 |
| SFTPC | P11686 | -1,57 | 0,000111636 |
| TNS1 | Q9HBL0 | -1,57 | 5,28E-06 |
| AEBP1 | Q8IUX7 | -1,58 | 6,68E-06 |
| C1orf198 | Q9H425 | -1,58 | 6,54E-05 |
| DMBT1 | Q9UGM3 | -1,58 | 0,026095359 |
| DMTN | Q08495 | -1,58 | 0,005592134 |
| TPPP | O94811 | -1,58 | 1,36E-05 |
| EHD1 | Q9H4M9 | -1,59 | 1,25E-05 |
| GNG2 | P59768 | -1,6 | 9,43E-09 |
| AKAP12 | Q02952 | -1,61 | 0,000432197 |
| ABCA8 | O94911 | -1,63 | 0,000505073 |
| EPB41L2 | O43491 | -1,63 | 3,30E-06 |

| | | | |
|---|---|---|---|
| CAMP | P49913 | -1,64 | 0,012253123 |
| HBG1 | P69891 | -1,64 | 0,0013919 |
| LIMS4 | P0CW20 | -1,64 | 1,04E-05 |
| SPTBN1 | Q01082 | -1,65 | 1,08E-06 |
| HTATIP2 | Q9BUP3 | -1,66 | 4,06E-05 |
| NHERF2 | Q15599 | -1,66 | 1,86E-06 |
| ITGA1 | P56199 | -1,68 | 7,37E-07 |
| RAB23 | Q9ULC3 | -1,68 | 3,47E-06 |
| PGM5 | Q15124 | -1,69 | 4,17E-05 |
| ANXA3 | P12429 | -1,7 | 1,86E-06 |
| SYNPO2 | Q9UMS6 | -1,7 | 0,004045203 |
| CNRIP1 | Q96F85 | -1,71 | 2,55E-06 |
| DNAH5 | Q8TE73 | -1,72 | 0,003308738 |
| PHF5A | Q7RTV0 | -1,72 | 7,37E-07 |
| DOCK9 | Q9BZ29 | -1,74 | 0,000260439 |
| APOC3 | P02656 | -1,75 | 0,000111559 |
| MGLL | Q99685 | -1,75 | 0,003825763 |
| SFTPA1 | Q8IWL2 | -1,77 | 0,00083151 |
| CLIC5 | Q9NZA1 | -1,79 | 2,30E-07 |
| CSMD2 | Q7Z408 | -1,79 | 0,013034107 |
| CYBRD1 | Q53TN4 | -1,79 | 5,35E-05 |
| PECAM1 | P16284 | -1,79 | 5,32E-07 |
| PF4 | P02776 | -1,8 | 0,006653969 |
| LRRK2 | Q5S007 | -1,81 | 0,00102395 |
| OGN | P20774 | -1,81 | 0,00283027 |
| JAM2 | P57087 | -1,84 | 5,87E-05 |
| PALM2AKAP2 | Q9Y2D5 | -1,84 | 2,84E-06 |
| ITGA2B | P08514 | -1,85 | 0,005976165 |
| SORBS3 | O60504 | -1,85 | 3,49E-06 |
| TNS2 | Q63HR2 | -1,85 | 3,49E-06 |
| ADD2 | P35612 | -1,86 | 0,000576932 |
| GYPC | P04921 | -1,86 | 8,33E-06 |
| UACA | Q9BZF9 | -1,86 | 4,12E-06 |
| WAS | P42768 | -1,86 | 5,53E-05 |
| GSTM5 | P46439 | -1,87 | 0,00238295 |
| NRP1 | O14786 | -1,87 | 3,17E-05 |
| SOD3 | P08294 | -1,87 | 9,82E-05 |
| MCAM | P43121 | -1,88 | 7,19E-05 |
| CA2 | P00918 | -1,9 | 3,02E-05 |
| CDH5 | P33151 | -1,9 | 2,30E-07 |
| ABI3BP | Q7Z7G0 | -1,91 | 0,00073793 |
| CHIT1 | Q13231 | -1,92 | 8,54E-06 |
| APOC2 | P02655 | -1,93 | 0,000401751 |
| CLDN5 | O00501 | -1,94 | 2,07E-07 |
| EVI2B | P34910 | -1,94 | 0,000158007 |

| | | | |
|---|---|---|---|
| GIMAP8 | Q8ND71 | -1,94 | 2,37E-06 |
| CD34 | P28906 | -1,96 | 4,50E-06 |
| EML1 | O00423 | -1,98 | 3,57E-05 |
| HBG2 | P69892 | -1,98 | 0,008958947 |
| IL33 | O95760 | -1,98 | 2,85E-05 |
| HBE1 | P02100 | -1,99 | 7,90E-05 |
| ITGA8 | P53708 | -1,99 | 0,000127459 |
| SIPA1L2 | Q9P2F8 | -1,99 | 0,000141513 |
| AK7 | Q96M32 | -2,02 | 0,000806452 |
| CA1 | P00915 | -2,02 | 0,000161776 |
| UBE2Q2 | Q8WVN8 | -2,02 | 0,000530868 |
| TPPP3 | Q9BW30 | -2,03 | 0,000549882 |
| OLFML3 | Q9NRN5 | -2,06 | 1,70E-05 |
| HBD | P02042 | -2,08 | 8,04E-05 |
| EHD2 | Q9NZN4 | -2,09 | 5,32E-07 |
| CAVIN2 | O95810 | -2,1 | 7,65E-08 |
| HSPA2 | P54652 | -2,12 | 6,70E-06 |
| CLEC14A | Q86T13 | -2,13 | 2,77E-05 |
| MCM5 | P33992 | -2,13 | 0,000232456 |
| CAVIN3 | Q969G5 | -2,15 | 1,92E-05 |
| HBB | P68871 | -2,15 | 3,25E-05 |
| KANK3 | Q6NY19 | -2,15 | 2,67E-05 |
| ANK1 | P16157 | -2,16 | 0,000588228 |
| CAMK1D | Q8IU85 | -2,17 | 1,59E-05 |
| HBA2 | P69905 | -2,17 | 3,38E-05 |
| RASIP1 | Q5U651 | -2,17 | 2,30E-07 |
| PODXL | O00592 | -2,18 | 2,37E-06 |
| SLC4A1 | P02730 | -2,18 | 0,000236869 |
| ACADL | P28330 | -2,2 | 7,82E-06 |
| STXBP6 | Q8NFX7 | -2,2 | 0,000116155 |
| ADIRF | Q15847 | -2,21 | 0,000115039 |
| RPS8 | P62241 | -2,22 | 1,81E-06 |
| CSTF2 | P33240 | -2,23 | 2,73E-07 |
| SLCO2A1 | Q92959 | -2,23 | 9,86E-05 |
| RHAG | Q02094 | -2,24 | 0,00014121 |
| CLIC3 | O95833 | -2,27 | 2,21E-06 |
| MME | P08473 | -2,28 | 0,000197634 |
| PRX | Q9BXM0 | -2,29 | 4,17E-05 |
| PDLIM2 | Q96JY6 | -2,3 | 5,12E-06 |
| AQP4 | P55087 | -2,32 | 1,61E-05 |
| CAVIN1 | Q6NZI2 | -2,32 | 4,00E-07 |
| EPB42 | P16452 | -2,34 | 1,59E-05 |
| ADIPOQ | Q15848 | -2,38 | 3,39E-05 |
| CDH13 | P55290 | -2,39 | 0,000137743 |
| CA4 | P22748 | -2,4 | 2,85E-05 |

| | | | |
|---|---|---|---|
| FHL1 | Q13642 | -2,4 | 2,04E-06 |
| ZW10 | O43264 | -2,45 | 1,06E-05 |
| SUSD2 | Q9UGT4 | -2,46 | 1,25E-05 |
| LIMCH1 | Q9UPQ0 | -2,47 | 7,37E-07 |
| SPARCL1 | Q14515 | -2,47 | 8,42E-06 |
| AOC3 | Q16853 | -2,48 | 4,00E-07 |
| CAV2 | P51636 | -2,52 | 7,66E-06 |
| GPRC5A | Q8NFJ5 | -2,53 | 2,73E-07 |
| HSPB6 | O14558 | -2,53 | 2,34E-05 |
| SCEL | O95171 | -2,54 | 2,13E-05 |
| ADH1B | P00325 | -2,57 | 3,74E-06 |
| UBR4 | Q5T4S7 | -2,59 | 1,25E-05 |
| LYVE1 | Q9Y5Y7 | -2,6 | 7,88E-06 |
| ACE | P12821 | -2,63 | 4,57E-05 |
| CD36 | P16671 | -2,68 | 2,53E-05 |
| ENPEP | Q07075 | -2,7 | 1,98E-05 |
| CAV1 | Q03135 | -2,78 | 6,40E-07 |
| ESAM | Q96AP7 | -2,81 | 6,89E-07 |
| OLFML1 | Q6UWY5 | -2,85 | 8,26E-06 |
| FABP4 | P15090 | -3,33 | 1,37E-05 |
| AGER | Q15109 | -5,22 | 1,26E-06 |

**Supplementary Table 2.** Deregulated proteins in extracellular vesicles (EVs) purified from Tumor-draining vein (TDV) as compared to EVs purified from Peripheral vein (PV) identified by mass spectrometry, according to the following thresholds: |log2 fold change| ≥ 1 and adjusted p value (Benjamini-Hochberg multiple testing correction) ≤ 0.05.

| Official Gene Symbol | Uniprot ID | $log_2$ Fold Change (TDV vs PV) | adjusted p value |
|---|---|---|---|
| MUC5B | Q9HC84 | 5,04 | 0,005163828 |
| CCT6A | P40227 | 3,74 | 0,00011637 |
| UNC45A | Q9H3U1 | 3,53 | 0,005163828 |
| IGHV5-51 | A0A0C4DH38 | 3,52 | 0,010761768 |
| ATP1A1 | P05023 | 3,46 | 0,001897142 |
| STXBP3 | O00186 | 3,46 | 0,00272099 |
| SSC5D | A1L4H1 | 3,28 | 0,000374093 |
| VDAC1 | P21796 | 3,28 | 0,000752903 |
| F12 | P00748 | 3,16 | 6,88E-06 |
| SRPRB | Q9Y5M8 | 3,15 | 0,001117865 |
| PRG4 | Q92954 | 3,08 | 1,75E-07 |
| TACC1 | O75410 | 3,06 | 0,009133315 |
| ECM1 | Q16610 | 3,03 | 9,84E-05 |
| ESYT1 | Q9BSJ8 | 2,99 | 0,000687073 |
| GPLD1 | P80108 | 2,87 | 0,003160593 |
| KLC1 | Q07866 | 2,82 | 0,000427937 |
| ANO10 | Q9NW15 | 2,81 | 0,004454711 |
| DOK3 | Q7L591 | 2,8 | 0,006977266 |
| PDLIM5 | Q96HC4 | 2,79 | 0,001398696 |
| ETHE1 | O95571 | 2,77 | 0,001419739 |
| CDSN | Q15517 | 2,73 | 0,000197884 |
| FBLN1 | P23142 | 2,72 | 0,000117672 |
| IGHV1-24 | A0A0C4DH33 | 2,68 | 0,010761768 |
| CSRP1 | P21291 | 2,63 | 0,013271342 |
| IST1 | P53990 | 2,62 | 0,005529207 |
| PSMA7 | O14818 | 2,61 | 0,005529207 |
| USP9X | Q93008 | 2,59 | 0,000427937 |
| TSPAN9 | O75954 | 2,58 | 0,002819229 |
| ATP8A1 | Q9Y2Q0 | 2,57 | 0,000241783 |
| SAA4 | P35542 | 2,47 | 0,000160054 |
| SPCS2 | Q15005 | 2,47 | 0,003160593 |
| KLKB1 | P03952 | 2,46 | 0,000193921 |
| VTN | P04004 | 2,46 | 4,30E-07 |
| SQOR | Q9Y6N5 | 2,45 | 0,001117865 |
| C1QC | P02747 | 2,43 | 7,07E-05 |
| GARS1 | P41250 | 2,41 | 1,56E-05 |
| IGHV4OR15-8 | A0A075B7B6 | 2,4 | 0,004749894 |
| IGHV4-28 | A0A0C4DH34 | 2,35 | 0,00464317 |
| FN1 | P02751 | 2,34 | 0,000427937 |

| | | | |
|---|---|---|---|
| DOCK8 | Q8NF50 | 2,33 | 0,00113822 |
| C1QA | P02745 | 2,31 | 0,000114547 |
| C1QB | P02746 | 2,3 | 0,00010095 |
| CHCHD3 | Q9NX63 | 2,3 | 0,003092678 |
| AK1 | P00568 | 2,28 | 0,012006362 |
| FCGR2A | P12318 | 2,27 | 0,00167845 |
| ITIH3 | Q06033 | 2,26 | 0,000922349 |
| GSR | P00390 | 2,25 | 0,025247146 |
| ITIH2 | P19823 | 2,24 | 0,001387605 |
| HADHA | P40939 | 2,22 | 0,000740997 |
| SERPINC1 | P01008 | 2,21 | 1,05E-05 |
| CPN1 | P15169 | 2,13 | 0,000160054 |
| SUCLG1 | P53597 | 2,08 | 8,69E-06 |
| MTHFD1 | P11586 | 2,05 | 0,005219907 |
| ATP5MK | Q96IX5 | 2,03 | 0,00860973 |
| MCEMP1 | Q8IX19 | 2,03 | 0,00161585 |
| UQCRC2 | P22695 | 2,03 | 0,004211563 |
| DBI | P07108 | 2,02 | 0,019083761 |
| NCKAP1 | Q9Y2A7 | 2,02 | 0,013819146 |
| LTF | P02788 | 1,99 | 0,010408772 |
| MYO9B | Q13459 | 1,98 | 0,000752903 |
| APOE | P02649 | 1,95 | 0,000752903 |
| C1S | P09871 | 1,95 | 0,000561361 |
| ROCK2 | O75116 | 1,94 | 0,003435149 |
| TAPBP | O15533 | 1,94 | 0,035816355 |
| LMAN2 | Q12907 | 1,93 | 0,003092678 |
| COMT | P21964 | 1,9 | 0,01047747 |
| GPD2 | P43304 | 1,9 | 0,003732614 |
| CFP | P27918 | 1,88 | 0,00113822 |
| MVP | Q14764 | 1,88 | 0,00113822 |
| NDUFS3 | O75489 | 1,87 | 0,00666639 |
| APOL1 | O14791 | 1,86 | 0,003160593 |
| PIGR | P01833 | 1,85 | 0,005163828 |
| ITIH1 | P19827 | 1,84 | 0,000193061 |
| IGHV3OR15-7 | A0A075B7D8 | 1,83 | 0,003435149 |
| APOC4 | P55056 | 1,82 | 0,010408772 |
| ERP29 | P30040 | 1,8 | 0,009357291 |
| GIT1 | Q9Y2X7 | 1,8 | 0,000632493 |
| APOA5 | Q6Q788 | 1,79 | 0,005274248 |
| PRAF2 | O60831 | 1,77 | 0,012869776 |
| AFG3L2 | Q9Y4W6 | 1,76 | 0,004211563 |
| AGPS | O00116 | 1,76 | 0,01752043 |
| CHMP6 | Q96FZ7 | 1,75 | 0,010527381 |
| GK | P32189 | 1,75 | 0,007439848 |
| PGLYRP2 | Q96PD5 | 1,75 | 0,027971085 |

| | | | |
|---|---|---|---|
| RTN1 | Q16799 | 1,75 | 0,007370395 |
| COL6A1 | P12109 | 1,73 | 0,010348489 |
| HRG | P04196 | 1,72 | 0,000460469 |
| MFGE8 | Q08431 | 1,72 | 0,010761768 |
| ANPEP | P15144 | 1,71 | 0,012869776 |
| ARL1 | P40616 | 1,71 | 0,000683682 |
| CHMP4A | Q9BY43 | 1,71 | 0,010761768 |
| APOB | P04114 | 1,7 | 0,005953852 |
| M6PR | P20645 | 1,7 | 0,000973656 |
| PZP | P20742 | 1,7 | 0,01207565 |
| SELPLG | Q14242 | 1,69 | 0,0072182 |
| VWA5A | O00534 | 1,68 | 0,019546533 |
| STIP1 | P31948 | 1,66 | 0,010761768 |
| GSTM3 | P21266 | 1,65 | 0,006882816 |
| PCYOX1 | Q9UHG3 | 1,63 | 0,005943017 |
| C1R | P00736 | 1,62 | 0,006977266 |
| AHNAK | Q09666 | 1,61 | 0,033453847 |
| BMP1 | P13497 | 1,61 | 0,002929196 |
| SLK | Q9H2G2 | 1,6 | 0,012735232 |
| TMEM40 | Q8WWA1 | 1,6 | 0,002161694 |
| SERPING1 | P05155 | 1,59 | 0,00011637 |
| UFM1 | P61960 | 1,58 | 0,024387411 |
| LRRC59 | Q96AG4 | 1,56 | 0,00860973 |
| HINT2 | Q9BX68 | 1,53 | 0,014347975 |
| NLRX1 | Q86UT6 | 1,53 | 0,016488416 |
| RHBDF2 | Q6PJF5 | 1,52 | 0,006057433 |
| EFR3A | Q14156 | 1,51 | 0,020097585 |
| GRHPR | Q9UBQ7 | 1,51 | 0,003619564 |
| PDGFB | P01127 | 1,51 | 0,005067556 |
| STX11 | O75558 | 1,49 | 0,003318568 |
| HSPD1 | P10809 | 1,48 | 0,005527433 |
| APOC1 | P02654 | 1,47 | 0,001909126 |
| ACOT9 | Q9Y305 | 1,46 | 0,023081524 |
| ALDH9A1 | P49189 | 1,44 | 0,019257889 |
| RTN4 | Q9NQC3 | 1,43 | 0,000427937 |
| CLTC | Q00610 | 1,42 | 0,0072182 |
| PLA2G7 | Q13093 | 1,42 | 0,013971599 |
| RAC2 | P15153 | 1,42 | 0,038398813 |
| CCT3 | P49368 | 1,4 | 0,001926899 |
| NAPG | Q99747 | 1,4 | 0,02224221 |
| NECTIN2 | Q92692 | 1,4 | 0,025251172 |
| NSF | P46459 | 1,4 | 0,012316773 |
| MENT | Q9BUN1 | 1,39 | 0,018292109 |
| C3 | P01024 | 1,38 | 0,000752903 |
| HLA-E | P13747 | 1,38 | 0,045583536 |

| | | | |
|---|---|---|---|
| MYO5A | Q9Y4I1 | 1,38 | 0,023007181 |
| PON1 | P27169 | 1,38 | 0,015104583 |
| TOMM70 | O94826 | 1,38 | 0,004465298 |
| DEFA1B | P59665 | 1,37 | 0,012869776 |
| IGKV1D-13 | A0A0B4J2D9 | 1,36 | 0,044358883 |
| DYNC1H1 | Q14204 | 1,35 | 0,003372625 |
| EML3 | Q32P44 | 1,35 | 0,00798054 |
| PDIA5 | Q14554 | 1,35 | 0,021780221 |
| CYB5R1 | Q9UHQ9 | 1,34 | 0,008125336 |
| KNG1 | P01042 | 1,33 | 0,00464317 |
| TM9SF3 | Q9HD45 | 1,33 | 0,022309222 |
| APOD | P05090 | 1,32 | 0,006284777 |
| APMAP | Q9HDC9 | 1,31 | 0,013470424 |
| HLA-A | P04439 | 1,3 | 0,045148108 |
| PSMC2 | P35998 | 1,27 | 0,010761768 |
| AMBP | P02760 | 1,26 | 0,005274248 |
| DCD | P81605 | 1,26 | 0,016995372 |
| FASN | P49327 | 1,26 | 0,023970281 |
| ATP5F1B | P06576 | 1,25 | 0,011741948 |
| ROCK1 | Q13464 | 1,25 | 0,014953809 |
| GGH | Q92820 | 1,24 | 0,011294657 |
| PGRMC1 | O00264 | 1,24 | 0,003445327 |
| COPS3 | Q9UNS2 | 1,23 | 0,043359024 |
| EXOC2 | Q96KP1 | 1,22 | 0,019811628 |
| ITPR2 | Q14571 | 1,17 | 0,024742561 |
| CANX | P27824 | 1,16 | 0,025247146 |
| CERS2 | Q96G23 | 1,15 | 0,015320762 |
| PI4KA | P42356 | 1,15 | 0,025247146 |
| IMMT | Q16891 | 1,14 | 0,004465298 |
| APOA2 | P02652 | 1,13 | 0,019167225 |
| GRB2 | P62993 | 1,13 | 0,020186629 |
| ANGPTL6 | Q8NI99 | 1,11 | 0,010527381 |
| CFH | P08603 | 1,11 | 0,047992194 |
| HSD17B10 | Q99714 | 1,11 | 0,025247146 |
| UBE2L3 | P68036 | 1,11 | 0,004601418 |
| CLINT1 | Q14677 | 1,1 | 0,035371893 |
| DSP | P15924 | 1,1 | 0,023237131 |
| DYNC1LI1 | Q9Y6G9 | 1,1 | 0,017346644 |
| KIF2A | O00139 | 1,1 | 0,048185135 |
| RTN2 | O75298 | 1,09 | 0,034705416 |
| CKAP5 | Q14008 | 1,08 | 0,023007181 |
| PLAA | Q9Y263 | 1,08 | 0,030698379 |
| HABP2 | Q14520 | 1,06 | 0,012869776 |
| IVD | P26440 | 1,06 | 0,025247146 |
| MPST | P25325 | 1,02 | 0,008166559 |

| | | | |
|---|---|---|---|
| F11 | P03951 | 1,01 | 0,0072182 |
| MTSS1 | O43312 | 1,01 | 0,025309349 |
| STXBP5 | Q5T5C0 | -1 | 0,033883098 |
| FTH1 | P02794 | -1,01 | 0,045148108 |
| HSPB1 | P04792 | -1,01 | 0,043359024 |
| PKM | P14618 | -1,01 | 0,012528461 |
| PSMB3 | P49720 | -1,01 | 0,020345051 |
| FYB1 | O15117 | -1,02 | 0,0072182 |
| GSTO1 | P78417 | -1,02 | 0,0492201 |
| TPM3 | P06753 | -1,03 | 0,020792797 |
| PDIA3 | P30101 | -1,04 | 0,007037555 |
| CFB | P00751 | -1,05 | 0,009827664 |
| CORO1A | P31146 | -1,05 | 0,010348489 |
| LYN | P07948 | -1,06 | 0,010239133 |
| SPN | P16150 | -1,07 | 0,016512733 |
| ALOX12 | P18054 | -1,08 | 0,024742561 |
| KALRN | O60229 | -1,08 | 0,047015725 |
| SPARC | P09486 | -1,08 | 0,03756841 |
| NAPA | P54920 | -1,1 | 0,025128912 |
| XPNPEP1 | Q9NQW7 | -1,1 | 0,034391058 |
| GNB2 | P62879 | -1,11 | 0,045103393 |
| MAPRE2 | Q15555 | -1,11 | 0,011212337 |
| PDE5A | O76074 | -1,13 | 0,015304526 |
| CD69 | Q07108 | -1,14 | 0,017711253 |
| TPT1 | P13693 | -1,14 | 0,02950035 |
| ACTR3 | P61158 | -1,15 | 0,002348986 |
| DAB2 | P98082 | -1,15 | 0,021780221 |
| GPSM3 | Q9Y4H4 | -1,15 | 0,020097585 |
| NIPSNAP3A | Q9UFN0 | -1,15 | 0,018459208 |
| MIF | P14174 | -1,16 | 0,005953852 |
| RAP2B | P61225 | -1,16 | 0,030698379 |
| ORM1 | P02763 | -1,17 | 0,017640205 |
| STEAP3 | Q658P3 | -1,17 | 0,04841568 |
| WASF2 | Q9Y6W5 | -1,17 | 0,038930642 |
| CD55 | P08174 | -1,18 | 0,016116212 |
| PGD | P52209 | -1,18 | 0,009873924 |
| PSMB2 | P49721 | -1,18 | 0,022408114 |
| GMFG | O60234 | -1,19 | 0,010348489 |
| PSMD2 | Q13200 | -1,2 | 0,038216134 |
| UBE2N | P61088 | -1,2 | 0,005953852 |
| VASP | P50552 | -1,2 | 0,006158341 |
| ADAM10 | O14672 | -1,21 | 0,046124199 |
| TSG101 | Q99816 | -1,21 | 0,010772917 |
| CAPZA2 | P47755 | -1,22 | 0,031687175 |
| CLIC4 | Q9Y696 | -1,22 | 0,009311764 |

| | | | |
|---|---|---|---|
| GP1BA | P07359 | -1,23 | 0,041851392 |
| MGLL | Q99685 | -1,23 | 0,007375681 |
| PRKCQ | Q04759 | -1,23 | 0,026408051 |
| ANXA5 | P08758 | -1,24 | 0,019201245 |
| CASP6 | P55212 | -1,24 | 0,049056759 |
| OTUB1 | Q96FW1 | -1,25 | 0,003092678 |
| PSMA1 | P25786 | -1,25 | 0,009188779 |
| SAR1A | Q9NR31 | -1,26 | 0,010761768 |
| RHEB | Q15382 | -1,28 | 0,041852867 |
| PPP1CA | P62136 | -1,29 | 0,00745538 |
| PRDX4 | Q13162 | -1,29 | 0,006158341 |
| DMTN | Q08495 | -1,3 | 0,011294657 |
| INPP4B | O15327 | -1,3 | 0,038930642 |
| TMED8 | Q6PL24 | -1,3 | 0,02676502 |
| PLEK | P08567 | -1,31 | 0,01239374 |
| LASP1 | Q14847 | -1,32 | 0,045148108 |
| PSMD13 | Q9UNM6 | -1,33 | 0,009188779 |
| EHD3 | Q9NZN3 | -1,34 | 0,00860973 |
| VTA1 | Q9NP79 | -1,34 | 0,036145489 |
| DNAJB6 | O75190 | -1,35 | 0,032237756 |
| ERBIN | Q96RT1 | -1,35 | 0,014977213 |
| ACTB | P60709 | -1,36 | 0,007439848 |
| CAPZB | P47756 | -1,37 | 0,005060358 |
| RAP1GDS1 | P52306 | -1,37 | 0,01467737 |
| HLA-DRA | P01903 | -1,38 | 0,013246652 |
| USP5 | P45974 | -1,38 | 0,011294657 |
| CAPNS1 | P04632 | -1,39 | 0,011221589 |
| TKT | P29401 | -1,39 | 0,02647131 |
| LDHB | P07195 | -1,4 | 0,008800811 |
| GP1BB | P13224 | -1,41 | 0,014461113 |
| PSMA3 | P25788 | -1,42 | 0,005802765 |
| LAT | O43561 | -1,43 | 0,008085796 |
| GP5 | P40197 | -1,44 | 0,005953852 |
| PTP4A2 | Q12974 | -1,45 | 0,003435149 |
| TAGLN2 | P37802 | -1,45 | 0,001427344 |
| ACTR1A | P61163 | -1,46 | 0,004535614 |
| POTEJ | P0CG39 | -1,46 | 0,013011404 |
| PPP2R1A | P30153 | -1,48 | 0,01373111 |
| DUSP3 | P51452 | -1,49 | 0,00161585 |
| GP6 | Q9HCN6 | -1,53 | 0,014519195 |
| GSTP1 | P09211 | -1,53 | 0,021241616 |
| YWHAZ | P63104 | -1,54 | 0,010527381 |
| DOK2 | O60496 | -1,55 | 0,003543991 |
| HSPA5 | P11021 | -1,55 | 0,001133384 |
| P2RX1 | P51575 | -1,55 | 0,011221589 |

| | | | |
|---|---|---|---|
| RAP1A | P62834 | -1,55 | 0,006158341 |
| GNAQ | P50148 | -1,56 | 0,022062286 |
| NAP1L1 | P55209 | -1,56 | 0,005060358 |
| ACTA1 | P68133 | -1,57 | 0,012530983 |
| DNM1L | O00429 | -1,57 | 0,000752903 |
| NCK2 | O43639 | -1,57 | 0,002161694 |
| PAK2 | Q13177 | -1,58 | 0,00745538 |
| PRKACB | P22694 | -1,58 | 0,020792797 |
| ACTR2 | P61160 | -1,59 | 0,006158341 |
| CLEC1B | Q9P126 | -1,6 | 0,038697599 |
| PARK7 | Q99497 | -1,6 | 0,001918819 |
| PRDX6 | P30041 | -1,6 | 0,013690512 |
| HIBCH | Q6NVY1 | -1,61 | 0,007690967 |
| PDIA4 | P13667 | -1,61 | 0,009928527 |
| S100A6 | P06703 | -1,61 | 0,001588033 |
| ATP6V0D1 | P61421 | -1,62 | 0,003697545 |
| HPCAL1 | P37235 | -1,63 | 0,007309002 |
| SLFN14 | P0C7P3 | -1,63 | 0,017752332 |
| HCLS1 | P14317 | -1,64 | 0,028352678 |
| S100A4 | P26447 | -1,64 | 0,006995953 |
| GC | P02774 | -1,65 | 0,000555018 |
| PLCB2 | Q00722 | -1,65 | 0,032904145 |
| TUBB4A | P04350 | -1,65 | 0,022024915 |
| XPO7 | Q9UIA9 | -1,65 | 0,035816355 |
| PDCD10 | Q9BUL8 | -1,67 | 0,005274248 |
| YWHAH | Q04917 | -1,67 | 0,014097472 |
| ARF3 | P61204 | -1,68 | 0,010761768 |
| ARHGDIB | P52566 | -1,68 | 0,004384462 |
| RNH1 | P13489 | -1,68 | 0,027271976 |
| KPNB1 | Q14974 | -1,69 | 0,004651889 |
| PRDX5 | P30044 | -1,69 | 0,017321264 |
| TPM4 | P67936 | -1,71 | 0,005953852 |
| TWF2 | Q6IBS0 | -1,71 | 0,016599066 |
| CAB39 | Q9Y376 | -1,72 | 0,000303396 |
| RANBP1 | P43487 | -1,72 | 0,017840008 |
| RSU1 | Q15404 | -1,72 | 0,003318568 |
| CSK | P41240 | -1,73 | 0,011636889 |
| ASL | P04424 | -1,75 | 0,048185135 |
| PPCS | Q9HAB8 | -1,75 | 0,034650694 |
| PSMB9 | P28065 | -1,75 | 0,009561612 |
| ARPC4 | P59998 | -1,77 | 0,000958604 |
| B2M | P61769 | -1,77 | 0,001419739 |
| ARPC2 | O15144 | -1,78 | 0,003076856 |
| LGALSL | Q3ZCW2 | -1,79 | 0,001419739 |
| SOD1 | P00441 | -1,79 | 0,00194803 |

| | | | |
|---|---|---|---|
| DDAH2 | O95865 | -1,8 | 0,001812206 |
| ILK | Q13418 | -1,8 | 0,003543991 |
| CFL1 | P23528 | -1,81 | 0,002389777 |
| USP14 | P54578 | -1,82 | 0,002739158 |
| CRLF3 | Q8IUI8 | -1,84 | 0,045799131 |
| MPP1 | Q00013 | -1,84 | 0,006158341 |
| STK38 | Q15208 | -1,84 | 0,000683682 |
| ATIC | P31939 | -1,86 | 0,008085796 |
| TMOD3 | Q9NYL9 | -1,86 | 0,003092678 |
| ARPC3 | O15145 | -1,87 | 0,002161694 |
| PPIA | P62937 | -1,87 | 0,001387605 |
| APOH | P02749 | -1,88 | 0,00011151 |
| PNP | P00491 | -1,88 | 0,002098122 |
| WDR1 | O75083 | -1,88 | 0,001419739 |
| TNIK | Q9UKE5 | -1,89 | 0,004214604 |
| GSPT1 | P15170 | -1,9 | 0,00478922 |
| COTL1 | Q14019 | -1,91 | 0,000958604 |
| DERA | Q9Y315 | -1,91 | 0,000722663 |
| ENO1 | P06733 | -1,91 | 0,001481559 |
| SEPTIN2 | Q15019 | -1,91 | 0,00113822 |
| PFN1 | P07737 | -1,92 | 0,002370816 |
| PGK1 | P00558 | -1,93 | 0,000752903 |
| SEPTIN7 | Q16181 | -1,94 | 0,001557169 |
| SH3BGRL3 | Q9H299 | -1,94 | 0,00206868 |
| GP9 | P14770 | -2 | 0,005943017 |
| CAPZA1 | P52907 | -2,02 | 0,002635819 |
| PPIF | P30405 | -2,09 | 0,023456819 |
| SARS1 | P49591 | -2,1 | 0,001059404 |
| CALD1 | Q05682 | -2,11 | 0,00021558 |
| ARPC5 | O15511 | -2,12 | 0,004353439 |
| GLUD1 | P00367 | -2,12 | 0,005163828 |
| EIF5A | P63241 | -2,14 | 0,001780933 |
| ETFB | P38117 | -2,14 | 0,006809226 |
| YWHAQ | P27348 | -2,16 | 0,005274248 |
| HBD | P02042 | -2,18 | 0,011965692 |
| ABI1 | Q8IZP0 | -2,23 | 0,0072182 |
| PTPN1 | P18031 | -2,23 | 0,021241616 |
| SRC | P12931 | -2,23 | 0,000752903 |
| HSPE1 | P61604 | -2,24 | 0,025247146 |
| PF4 | P02776 | -2,25 | 0,000534311 |
| TPM1 | P09493 | -2,28 | 0,002431113 |
| GDI2 | P50395 | -2,32 | 0,019201245 |
| UBE2V1 | Q13404 | -2,34 | 0,011390851 |
| GET3 | O43681 | -2,36 | 0,022309222 |
| PTPRJ | Q12913 | -2,36 | 0,003127016 |

| | | | |
|---|---|---|---|
| TMBIM1 | Q969X1 | -2,45 | 0,022024915 |
| TOM1L2 | Q6ZVM7 | -2,46 | 0,014754364 |
| STX4 | Q12846 | -2,47 | 0,001909126 |
| UGP2 | Q16851 | -2,47 | 0,005045741 |
| YWHAB | P31946 | -2,49 | 0,000722663 |
| HSPA8 | P11142 | -2,52 | 0,000561361 |
| YWHAE | P62258 | -2,52 | 0,000752903 |
| ITGB2 | P05107 | -2,58 | 0,023007181 |
| HLA-B | P01889 | -2,7 | 0,000257986 |
| SCAMP2 | O15127 | -2,75 | 0,014519195 |
| IGHD | P01880 | -2,78 | 0,027819184 |
| PSME1 | Q06323 | -2,87 | 0,000174633 |
| LAP3 | P28838 | -3,69 | 0,000160054 |
| ATP2B4 | P23634 | -3,73 | 0,007375681 |
| MYH7B | A7E2Y1 | -4,28 | 0,010400085 |
